# A genome-scale atlas reveals complex interplay of transcription and translation in an archaeon

**DOI:** 10.1101/2022.08.31.505529

**Authors:** Alan P. R. Lorenzetti, Ulrike Kusebauch, Lívia S. Zaramela, Wei-Ju Wu, João P. P. de Almeida, Serdar Turkarslan, Adrián L. G. de Lomana, José V. Gomes-Filho, Ricardo Z. N. Vêncio, Robert L. Moritz, Tie Koide, Nitin S. Baliga

**Affiliations:** Department of Biochemistry and Immunology, Ribeirão Preto Medical School, University of São Paulo, Ribeirão Preto, Brazil; Institute for Systems Biology, Seattle, WA, USA; Institute of Biological Sciences, Federal University of Minas Gerais, Belo Horizonte, Brazil; Center for Systems Biology, University of Iceland, Reykjavik, Iceland; Prokaryotic RNA Biology, Phillips-Universität Marburg, Marburg, Germany; Department of Computation and Mathematics, Faculty of Philosophy, Sciences and Letters at Ribeirão Preto, University of São Paulo, Ribeirão Preto, Brazil; Departments of Biology and Microbiology, University of Washington, Seattle, WA, USA; Molecular and Cellular Biology Program, University of Washington, Seattle, WA, USA; Lawrence Berkeley National Lab, Berkeley, CA, USA

## Abstract

The scale of post-transcriptional regulation and the implications of its interplay with other forms of regulation on environmental acclimation is underexplored for organisms of the domain Archaea. Here, we have investigated the scale of post-transcriptional regulation in the extremely halophilic archaeon *Halobacterium salinarum* NRC-1 by integrating transcriptome-wide locations of transcript processing sites (TPS) and SmAP1 binding, genome-wide locations of antisense RNAs (asRNAs), and consequences of RNase_2099C knockout on differential expression of all genes. This integrated analysis has discovered that 54% of all protein-coding genes in the genome of this haloarchaeon are likely targeted by multiple mechanisms for putative post-transcriptional processing and regulation, with about 20% of genes likely regulated by combinatorial schemes involving SmAP1, asRNAs, and RNase_2099C. Comparative analysis of mRNA levels (RNA-Seq) and protein levels (SWATH-MS) for 2,579 genes over four phases of batch culture growth in complex medium has generated additional evidence for conditional post-transcriptional regulation of 7% of all protein-coding genes. We demonstrate that post-transcriptional regulation may act to fine-tune specialized and rapid acclimation to stressful environments, e.g., as a switch to turn on gas vesicle biogenesis to promote vertical relocation in anoxic conditions and to modulate frequency of transposition by IS elements of the IS*200*/IS*605*, IS*4*, and IS*H3* families. Findings from this study are provided as an atlas in a public web resource (https://halodata.systemsbiology.net).

**IMPORTANCE:** While the transcriptional regulation landscape of archaea has been extensively investigated, we currently have limited knowledge about post-transcriptional regulation and its driving mechanisms in this domain of life. In this study, we collected and integrated omics data from multiple sources and technologies to infer post-transcriptionally regulated genes and the putative mechanisms modulating their expression at the protein level in *Halobacterium salinarum* NRC-1. The results suggest that post-transcriptional regulation may drive environmental acclimation by regulating hallmark biological processes. To foster discoveries by other research groups interested in the topic, we extended our integrated data to the public in the form of an interactive atlas (https://halodata.systemsbiology.net).

## INTRODUCTION

By virtue of their co-existence with multiple organisms within a community, microbes are under significant evolutionary selection pressure to maximize resource utilization for growth and sustenance, while minimizing waste (1). For this reason, even within their streamlined genomes, microbes possess extensive regulatory mechanisms at multiple levels of information processing (2–5). While regulation at the transcriptional level is typically modular with genome-wide consequences (4, 6), regulation at the post-transcriptional level is believed to be more nuanced and localized to specific sets of functions that are directly associated with environment-specific phenotypic traits (7). In other words, while transcriptional regulation mediates large-scale physiological adjustments, post-transcriptional regulation fine-tunes specific functions to optimize environmental acclimation. Understanding the interplay of regulation across the different layers of information processing will give insight into how microbes compete and collaborate effectively with other co-inhabiting organisms. In addition to having foundational significance, these insights also have important implications for synthetic biology approaches to introduce novel traits while minimizing fitness tradeoffs in an engineered organism (8–11).

Understanding the interplay of regulation across transcription and translation in organisms of the domain Archaea is especially interesting for several reasons. First, while they have been discovered across diverse environments, archaea are particularly known for specialized phenotypic adaptations for some of the most extreme and dynamic habitats (12). Second, archaea are unique in terms of possessing a mix of information processing mechanisms that are distinctly eukaryotic or bacterial. For instance, while their general transcriptional machinery including the RNA polymerase shares ancestry with eukaryotic counterparts, the regulation of transcription is mediated by regulators that have bacterial ancestry (13, 14). There has been extensive work across several archaeal model organisms that has characterized basal transcription and its regulation both in molecular detail and at a systems level (2, 3, 15). By contrast, it has been only in the recent past that we have begun to appreciate the role of post-transcriptional regulatory mechanisms in specialized phenotypic acclimation of archaea. There is evidence that translational efficiency in methanogenic archaea is modulated through differential processing of 5’ UTRs (16), mRNA secondary structures (17), or context-specific binding by small regulatory RNAs (sRNAs) to conditionally occlude ribosome binding sites within transcripts (18) or to stabilize them (19). Studies conducted in a psychrophile have discovered that post-transcriptional regulation directly influences methanol conversion into methane at lower temperatures (20). Similarly, RNase-mediated disruption of positive autoregulation of potassium uptake was discovered to be an important mechanism for energetically-efficient and rapid acclimation of a halophile in a salinity shift scenario (21). These examples illustrate how some archaea utilize post-transcriptional regulation to fine-tune specific functions and pathways for specialized phenotypic acclimation to environmental change.

However, a lot remains to be understood regarding the scale of post-transcriptional regulation in archaea and the extent to which they are deployed in combinatorial schemes to fine-tune phenotypes for environmental acclimation. For instance, the widely conserved and extensively characterized RNA-binding proteins (RBP), including Csp (A, C, and E), CsrA, RNaseE, YbeY, and Hfq, are known to play important post-transcriptional regulatory functions in bacteria (22), but there is limited understanding of the roles of their orthologs in archaea. Hfq is a member of an RNA-guided complex, a well-characterized bacterial RNA chaperone known to interfere in mRNA translation (23, 24), which acts in a manner analogous to the RNA-induced silencing complex (RISC) in eukaryotes to regulate specific mRNAs (25). Notably, the Hfq homolog, Sm-like archaeal protein (SmAP1 or Lsm), has been characterized structurally across multiple archaea (26–29), including *Halobacterium salinarum* NRC-1 (30), and shown to likely mediate post-transcriptional regulation through sRNA-binding in *Haloferax volcanii* (31, 32) and *Sulfolobus solfataricus* (33). However, we do not understand the mechanism, importance, context or scale of post-transcriptional regulation mediated by SmAP1 (and other RBPs) (34, 35) or, for that matter, by the large numbers of sRNAs, antisense RNAs (asRNAs), and RNases that have been discovered across archaeal genomes (36).

Here, we have investigated the scale of interplay between transcriptional and post-transcriptional mechanisms in regulating protein levels in the halophilic archaeon *H. salinarum* NRC-1, which has served as a model to investigate traits of organisms in the domain Archaea. In particular, *H. salinarum* NRC-1 has been widely used as a model organism to dissect hallmark traits of halophilic archaea, including niche adaptation via expanded families of general transcription factors (37), large-scale genome organization by genomic repeats and insertion sequences (IS) (38, 39), flotation by gas vesicle biogenesis (40), phototransduction by bacteriorhodopsin (41), and how modularity of translational complexes enables rapid acclimation to environmental changes (42). Prior work has characterized at a systems level and in mechanistic detail many aspects of the global transcriptional regulatory network of *H. salinarum* NRC-1 (2, 3), with extensive validations through genetic perturbation studies and physical mapping of genome-wide protein-DNA interactions of multiple transcription factors (4, 5). However, the transcriptional regulatory network by itself or the half-lives of all transcripts (43) did not fully explain the complex relationship between absolute and relative abundance of transcripts and proteins across different environmental contexts (44, 45), suggesting an important role for post-transcriptional regulation. Indeed, prior studies have uncovered evidence for the potential of extensive post-transcriptional regulation in *H. salinarum* NRC-1, including the presence of a strikingly large number of regulatory elements within coding sequences (3) that leads to widespread conditional splitting of at least 40% of all operons into multiple overlapping transcriptional units (5), presence of asRNAs for 22% of all genes (46), differential regulation of 23 transcripts in an RNase knockout background (21), and extensive transcript processing sites (TPS) across 43% of all coding sequences (47).

Through integrated analysis of a new transcriptome-wide map of SmAP1 binding located with RNA immunoprecipitation sequencing (RIP-Seq), global differential regulation of transcripts upon deletion of an RNase (VNG_2099C) implicated in acclimation to salinity change (21), and locations of asRNAs and TPS (46, 47), we have generated a genome-scale atlas that has discovered that 54% of all protein-coding genes in *H. salinarum* NRC-1 are targeted by multiple mechanisms for putative post-transcriptional regulation. Interestingly, 20% of all protein-coding genes are likely post-transcriptionally regulated in combinatorial schemes involving SmAP1, asRNAs, and RNase. Further, through comparative analysis of dynamic changes in mRNA levels (RNA-Seq), ribosome footprints (Ribo-Seq) (42), and protein levels (SWATH-MS) (Kusebauch et al., in preparation) for 2,579 representative genes over four phases of batch culture growth in complex medium, we have generated evidence that 7% of all protein-coding genes (188 genes) are indeed post-transcriptionally regulated. Notably, 78% of these post-transcriptionally regulated genes were mechanistically associated with SmAP1-binding, asRNAs, TPS, and/or RNase-mediated differential regulation. Through in-depth analysis we demonstrate how post-transcriptional regulation acts to fine-tune specialized environmental acclimation, e.g., as a switch to turn on gas vesicle biogenesis and to modulate frequency of transposition by IS elements of the IS*200*/IS*605*, IS*4*, and IS*H3* families. Finally, we have generated an interactive web resource to support collaborative community-wide exploration and characterization of the *H. salinarum* NRC-1 multi-omics Atlas (https://halodata.systemsbiology.net).

## RESULTS

### Evidence for post-transcriptional regulation by SmAP1, asRNAs, and RNase_2099C

Since the publication of its genome sequence in 2000, multiple sources of gene annotations have emerged for *H. salinarum* NRC-1 (48–50). To standardize annotations, we clustered sequences from each source to eliminate redundancy while differentiating between paralogs (see Methods; Table S1; File S1). In summary, this analysis identified 2,631 non-redundant transcripts, including 2,579 coding and 52 non-coding RNAs (rRNAs, tRNAs, signal recognition particle RNA, and RNase P) with a dictionary anchored by locus tags from (50) and mapped to locus tags of the closely related strain *H. salinarum* R1 (File S1).

Next, we compiled orthogonal, genome-wide evidence for putative post-transcriptional regulation. Specifically, we relocated one or more published transcript processing sites (TPS) within at least 966 protein-coding genes (37% of all protein-coding genes) (47), mapped cis-acting asRNAs for 536 genes (46), and determined that 166 genes were differentially expressed upon deletion of one out of 12 RNases predicted within the genome (VNG_2099C; here onwards “RNase_2099C”) (21) (File S2). To characterize the role of SmAP1 (VNG_1496G) in *H. salinarum* NRC-1, epitope-tagged SmAP1-RNA complexes were co-immunoprecipitated from late-exponential phase cultures from standard growth conditions (Figure S1), and transcriptome-wide binding locations of SmAP1 were mapped by enrichment of sequenced transcripts (RIP-Seq; see Methods). Consistent with previous *in vitro* observations from diverse archaea, the RIP-Seq analysis discovered that SmAP1 preferentially binds to AU-rich transcripts (Figure S2A) (28–31, 51). In particular, we determined that SmAP1 binds to 15% (397/2,579) of all protein-coding transcripts in *H. salinarum* NRC-1, including its own coding transcript (File S1), suggesting putative autoregulation in light of the observed dynamics for mRNA and protein levels (Figure S2B).

Integrated analysis of locations of SmAP1 binding, asRNAs, and TPS, and differential expression in Δ*RNase_2099C* revealed that at least 1,394 genes were potentially subject to post-transcriptional regulation by at least one of these mechanisms, with 514 genes under putative combinatorial regulation by two or more mechanisms (Figure 1). Interestingly, transcripts that were upregulated in the Δ*RNase_2099C* strain background were preferentially bound by SmAP1 (*p*-value = 0.02), associated with cognate asRNAs (*p*-value = 0.04), and enriched for TPS (*p*-value = 6.7×10^−5^). These findings could suggest that SmAP1 and asRNAs are responsible for the recruitment of RNase_2099C to mediate targeted cleavage of transcripts. Thus, the integrated analysis predicted that 20% to 54% of the *H. salinarum* genome is post-transcriptionally regulated (Figure 1; 514 to 1,394 out of 2,579 genes). The fact that SmAP1, asRNAs, and RNase_2099C account for putative regulation of 858 genes, suggests that myriad mechanisms, potentially involving other RBPs and RNases noted above, are likely at play even in the limited conditions represented in standard growth conditions.

**Figure 1.**
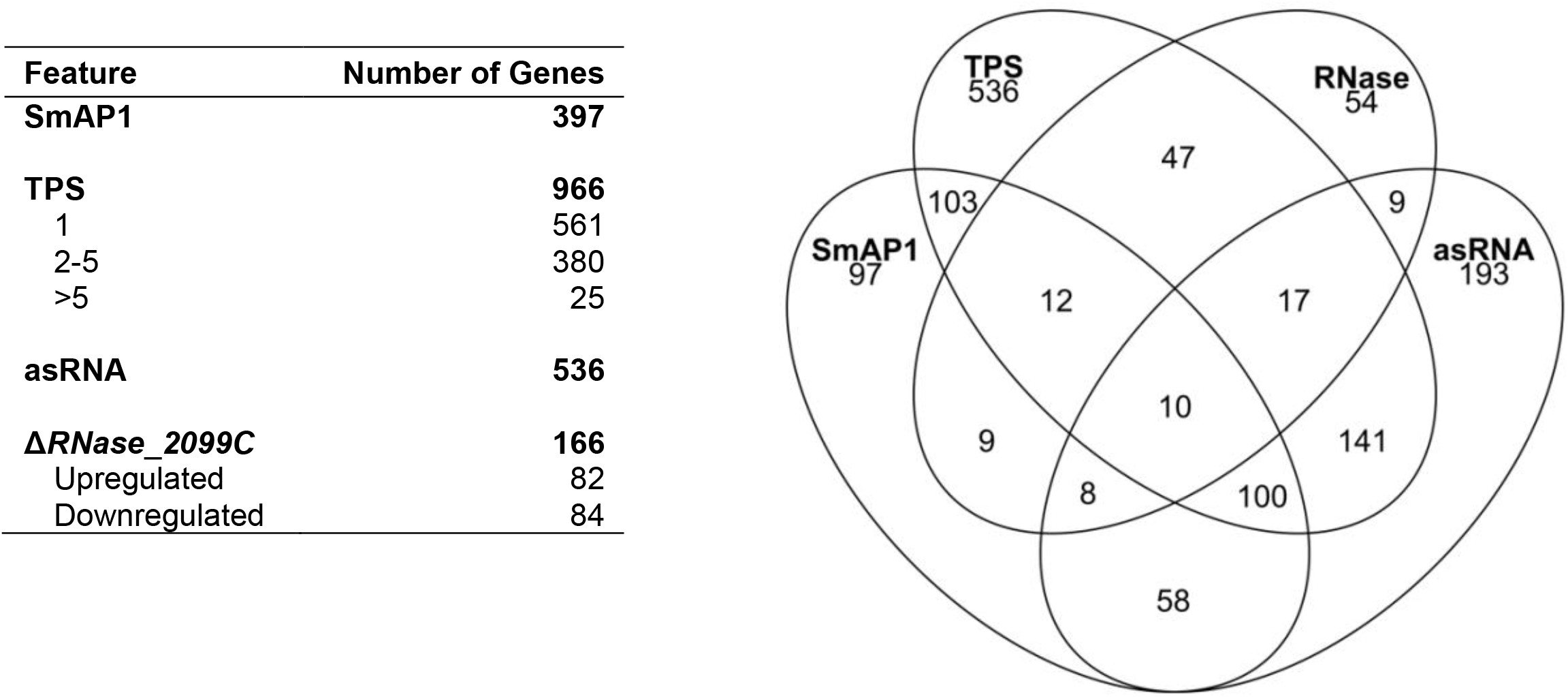
Features potentially associated with post-transcriptional regulation. Four features related to the post-transcriptional regulation in *H. salinarum*. Sets are comprised of genes that bind to SmAP1, show transcript processing sites (TPS), have a putative cis-regulatory antisense RNA (asRNA), and are differentially expressed in the RNase_2099C knockout strain (Δ*VNG_2099C*).

### Evidence of post-transcriptional regulation in global trends of mRNA and protein levels

We investigated concordance in patterns of absolute abundance at the transcriptional and translational levels for each gene by calculating Pearson correlation coefficients between mRNA and protein quantification across all the sampled physiological states (*R*_TP1_ = 0.67; *R*_TP2_ = 0.68; *R*_TP3_ = 0.57; *R*_TP4_ = 0.44) (Figure 2A-D). The weaker correlation (*R*_TP1_ = *R*_TP2_ > *R*_TP3_ > *R*_TP4_; Table S2) in later stages of batch culture growth was skewed towards repression of translation; that is, highly abundant mRNAs were associated with low abundance proteins in the quiescent physiological state (TP4). We also noticed that protein levels correlated slightly better with mRNA levels from the previous time point (*R*_P-TP2 m-TP1_ = 0.68; *R*_P-TP3 m-TP2_ = 0.67; *R*_P-TP4 m-TP3_ = 0.57; Figure 2E-G; Table S2), which is consistent with the sequential and temporal relationship between transcription and translation, as we have previously shown (44, 45). We discovered that 6.5% of all protein-coding genes (167) with high mRNA levels (upper quintile) were associated with low protein levels (lower quintile or undetected) over some or all four stages of growth in batch culture (Figure S3A, File S3). Specifically, the 167 genes were enriched for SmAP1 binding, asRNAs, and TPS (*p*-value = 2.3×10^−4^, 2.9×10^−2^, and 1.1×10^−7^, respectively) and had longer average mRNA half-lives (13.7 min. vs. 12.3 min.; *p*-value = 1.1×10^−2^). Within this set, 64 genes associated with protein levels detected in the lower quintile (green points in Figure 2A-D; Figure S3B; File S3) were enriched for TPS (*p*-value = 2.6×10^−4^). A second set of 117 genes, whose proteins were not detected likely due to their low levels or complete absence (see Methods; Figure S3C; File S3), was enriched for SmAP1 binding and TPS (*p*-value = 1.7×10^−6^ and 2.8×10^−6^, respectively), had longer average mRNA half-lives (14.2 min. vs. 12.3 min; *p*-value = 2.7×10^−3^), and was upregulated in Δ*RNase_2099C* strain (*p*-value = 1.5×10^−2^). Refer to File S4 for sets and tests.

**Figure 2.**
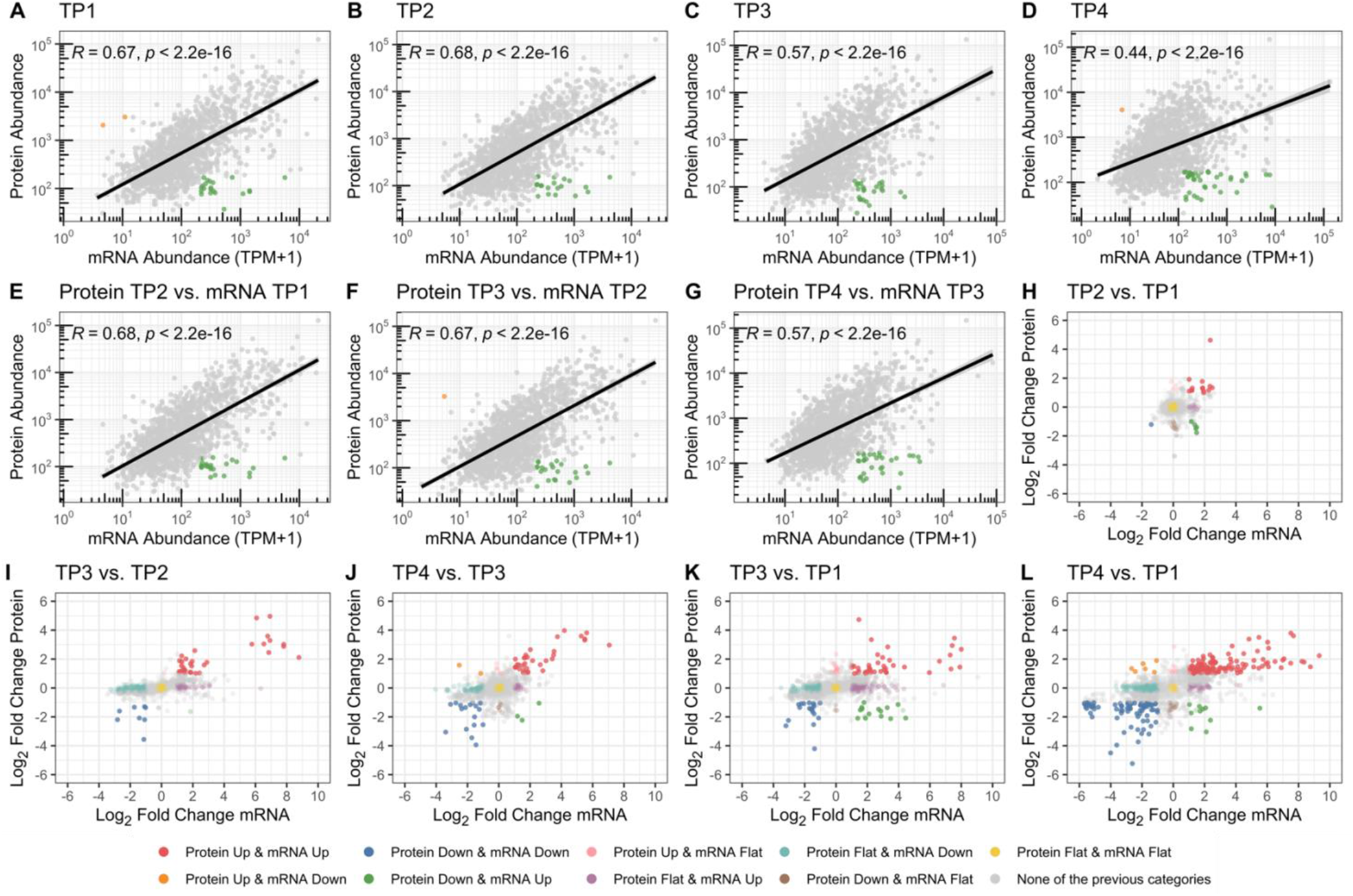
Genes following patterns compatible with post-transcriptional regulation. Each panel shows protein (*y*-axis) and mRNA (*x*-axis) absolute abundance (log_10_-transformed) or relative changes (log_2_ fold change). Absolute abundance-based analysis is reported from **A** to **D** in a time point-wise manner and from **E** to **G** in a time-lag perspective. Gray points represent entities following usual patterns; orange points represent entities within the upper quintile of protein abundance and lower quintile of mRNA abundance; green points represent entities within the lower quintile of protein abundance and upper quintile of mRNA abundance. The solid black line illustrates the fitted linear regression model. **H, I**, and **J** present the relative abundance-based analysis of protein and mRNA levels in consecutive physiological state transitions. **K** and **L** present the same variables for long physiological state transitions. Points are color-coded according to multiple combinations of change status considering both variables. TP1: early exponential growth phase; TP2: mid-exponential growth phase; TP3: late exponential growth phase; TP4: stationary phase.

Finally, we searched for potentially post-transcriptionally regulated genes by correlating dynamic relative changes in protein and mRNA levels over time (Figure 2H-L; File S5; File S6). For example, during the transition from TP1 to TP2, we observed a decrease in protein abundance of five transcriptionally upregulated genes over the same timeframe (Figure 2H). This cluster (Figure S4; File S6), comprised of five genes depicted by green points (VNG_7025, VNG_7026, VNG_7039, VNG_7103, and VNG_6313G) in Figure 2H, with enrichment for SmAP1 binding, asRNAs, and TPS (*p*-value = 8.5×10^−5^, 3.8×10^−4^, and 0, respectively), is a strong candidate for post-transcriptional repression. The genes also had lower codon adaptation index (CAI; 0.64 vs. 0.77; *p*-value = 3.9×10^−3^) and increased mRNA levels in the *ΔRNase_2099C* strain (log_2_ fold change = 1 vs. 0.02; *p*-value = 3.5×10^−4^). The comparative analysis of mRNA and protein abundance changes across all transition states (TP1 to TP2, TP2 to TP3, TP3 to TP4, TP1 to TP3 and TP1 to TP4) identified 26 potentially post-transcriptionally repressed transcripts (Figure S5; File S6) enriched for SmAP1 binding and TPS (*p*-value = 3.5×10^−3^ and 2.3×10^−4^, respectively), and upregulated in Δ*RNase_2099C* strain (*p*-value = 9.2×10^−7^). Again, refer to File S4 for sets and tests.

Altogether, the combined analyses of correlations between absolute and relative abundance of mRNAs and proteins provided further evidence for post-transcriptional regulation of at least 7% of all genes (188 out of 2,579) in *H. salinarum* NRC-1 during transition from active growth to the stationary phase. Notably, 78% of these genes (147/188) with poor mRNA-protein correlation were among the 1,394 genes associated with putative post-transcriptional regulation features, including SmAP1 binding, asRNAs, and TPS (*p-*value = 1.9×10^−9^, 7.6×10^−6^, and 2.5×10^−21^, respectively). Together these findings suggest complex combinatorial post-transcriptional regulation of these genes at specific growth stages.

### *Construction of the* H. salinarum *NRC-1 multi-omics Atlas*

To facilitate discovery of evidence of post-transcriptional regulation, we compiled corresponding quantitation of mRNAs (RNA-Seq), ribosome-protected mRNA fragments (RPF; Ribo-Seq) (42), and proteins (SWATH-MS) (Kusebauch et al., in preparation), quantile normalized them (File S1) for scale adjustment, and performed calculations of translational efficiency (TE) and ribosome occupancy (RO) for 2,579 genes across early exponential (TP1), mid-exponential (TP2), late-exponential (TP3), and stationary (TP4) phases of growth in batch culture (see Methods; Figure 3; File S7). Further, we also included general properties such as GC content, mRNA half-life, and CAI for each gene, as they are known to influence dynamics of the interplay between transcription and translation (43, 52) and could likely explain discrepant patterns of corresponding changes across mRNA, RPF, and proteins. A quick exploratory analysis of GC content and CAI, brought up their association to protein levels in this study (Figure S6). Genes in the atlas were organized into nine groups based on patterns of absolute abundance (File S3) and relative changes across mRNA and protein levels (File S6). This analysis revealed that at least 188 genes (7% of all protein-coding genes in the atlas) had incoherent mRNA-protein correlation patterns across the four physiological states during growth in batch culture. Notably, 147 of these 188 genes were associated with at least one post-transcriptional regulation mechanism noted above. The *H. salinarum* NRC-1 Atlas is accessible through an application (https://halodata.systemsbiology.net) that supports interactive exploration by zooming into specific segments of a heatmap, by searching for genes of interest, or through a searchable genome browser. The following sections demonstrate how the atlas facilitates in-depth investigations into post-transcriptional regulation of hallmark processes in *H. salinarum* NRC-1.

**Figure 3.**
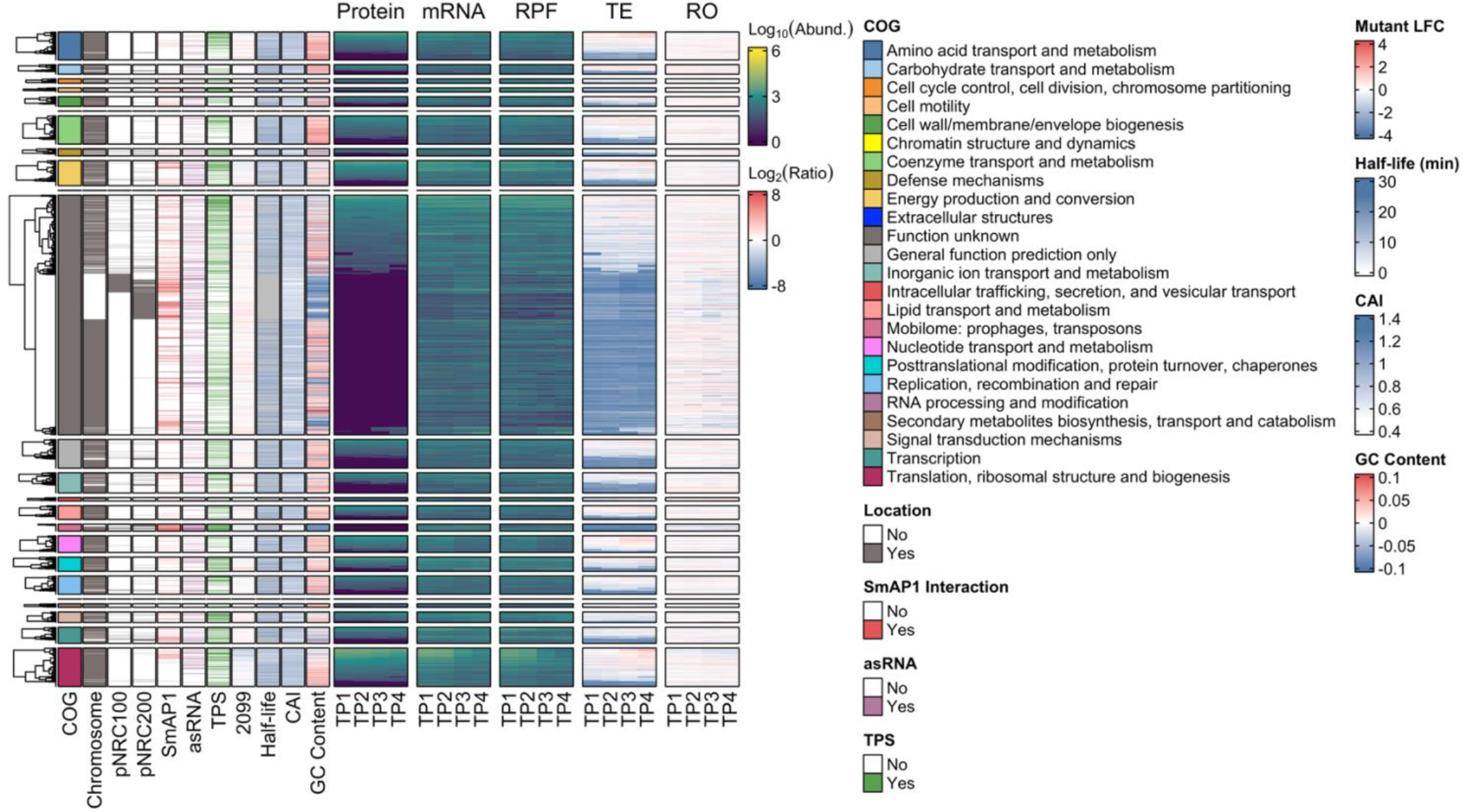
An atlas of the transcriptome, ribosome profile, and proteome for *Halobacterium salinarum* NRC-1. The heatmap shows quantile-normalized log_10_-transformed abundance levels for proteins (a pseudocount was imputed for missing values), messenger RNAs (mRNAs; TPM+1), and ribosome-protected mRNA fragments (RPF; TPM+1) for 2,579 genes across four consecutive stages of batch culture growth, namely early exponential, mid-exponential, late exponential, and stationary phase (TP1, TP2, TP3, and TP4, respectively). Log_2_-transformed translational efficiency (TE) and ribosome occupancy (RO) were computed by dividing protein levels by mRNA levels and mRNA levels by RPF levels, respectively. We present general features on the left-hand side, starting with the cluster of orthologous genes (COG) functional categories (97), split into groups before clustering the protein levels. Chromosome, pNRC100, and pNRC200 show the replicon location of each gene within the genome. The presence of SmAP1 binding, antisense RNAs (asRNA) (46), and putative endoribonuclease-generated transcript processing sites (TPS) (47) are indicated in corresponding tracks. The 2099 track shows log_2_ fold change (LFC) of transcript levels in the RNAse_2099C null mutant (Δ*VNG_2099C*) relative to the parent Δ*ura3* strain (21). mRNA half-lives (43), codon adaptation index (CAI), and the deviation of GC content from average GC content of all transcripts are also indicated in corresponding tracks. See inset keys for color codes for each track and Methods section for details. Interactive and expanded static versions of this figure are available in our *H. salinarum* NRC-1 multi-omics Atlas portal (https://halodata.systemsbiology.net).

### *Functional implications of growth-associated post-transcriptional regulation in* H. salinarum

Altogether, the comparison of absolute and relative abundance of mRNA and protein levels yielded evidence for post-transcriptional regulation of 188 genes during batch culture growth (Figure 2; File S3; File S6). Furthermore, the longer transcript half-lives together with enrichment of SmAP1-binding, asRNAs, TPS, and differential regulation upon deletion of RNase_2099C provided evidence for post-transcriptional processing, and associated putative mechanisms of regulation of different gene subsets. While a substantial number of genes were of unknown function, important processes were represented among genes of known functions; these included gas vesicle biogenesis, transposition-mediated genome reorganization, motility, translation, and energy transduction (Figure 4). Among these, both gas vesicles and extensive genome reorganization mediated by activity of mobile genetic elements are hallmark traits of *H. salinarum* NRC-1 that are triggered in specific environmental contexts, including late growth and stationary phases. Below, we present vignettes on each of these two processes to illustrate how the *H. salinarum* NRC-1 multi-omics Atlas enables the discovery of mechanistic insight into post-transcriptional regulation of specific phenotypes.

**Figure 4.**
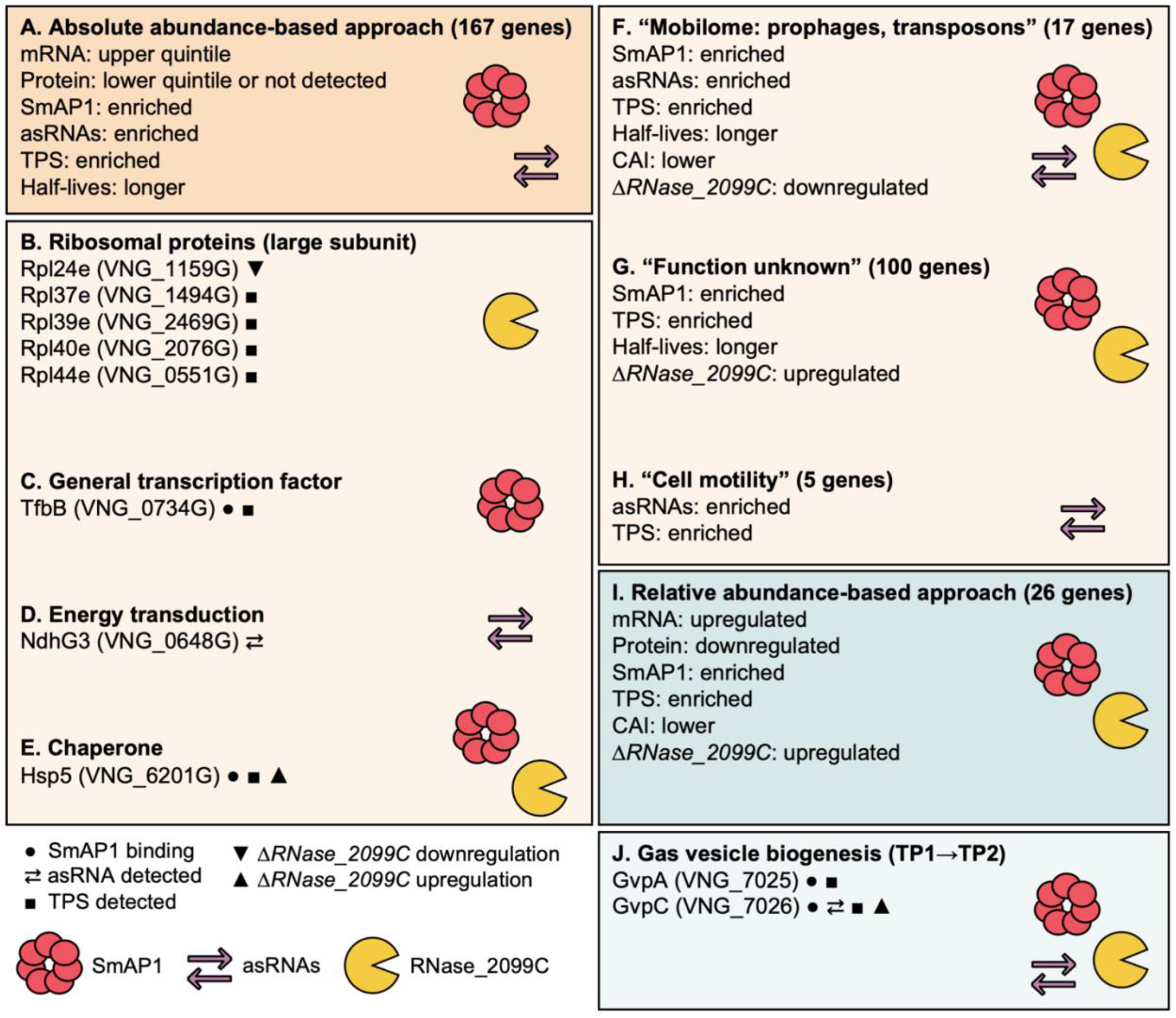
Functions of putative post-transcriptionally regulated genes and potential driving mechanisms. The figure shows the common properties of groups of putative post-transcriptionally regulated genes. **A**. The union set of genes found by the absolute abundance-based approach across the growth curve (green points in Figure 2A-D). **B-E**. Arbitrarily selected genes of known functions (subsets of **A**). **F-H**. Gene categories according to clusters of orthologous genes (COG) with enriched features compatible with the post-transcriptional regulation hypothesis (subsets of **A**). **I**. The union set of genes found by the relative abundance-based approach across the growth curve (upregulated mRNA and downregulated protein; green clusters in Figure 2H-L). **J**. Genes of the *gvp* cluster in the transition from early exponential (TP1) to mid-exponential growth phase (TP2) (subset of **I**). See File S4 for a complete list of genes within each group (**A, F-H, I**) and the respective supporting evidence. TPS: Transcript processing sites; asRNA: antisense RNA; CAI: Codon adaptation index.

#### The role of SmAP1 in the regulation of transposition and genome reorganization

Transposases are typically encoded within insertion sequences (IS), a type of transposable element that is ubiquitous across prokaryotes, and known to mediate self-mobilization to new locations in the genome (53, 54). *H. salinarum* NRC-1 mobilome is comprised by 80 full and 33 partial IS elements of eight families (ISfinder/ISbrowser) (55, 56), some of which are known to introduce phenotypic diversity in flotation, by disrupting *gvp* locus at 1-5% frequency, and also in phototrophic energy production, by disrupting the bacteriorhodopsin gene (*bop*) locus at 0.01% frequency, potentially driving niche acclimation in brine pools (38, 57, 58). Notably, SmAP1 bound 24 of the 33 mobilome transcripts (Figure 5A; Figure S2C; enrichment *p*-value = 10^−14^), consistent with their low GC content (Figure 5B) and the previously implicated role of its bacterial homolog in regulating transposition events (59, 60). Out of the 33 mobilome proteins, only four were detected at the protein level (Figure 5AC), including three TnpB proteins encoded by IS elements of the IS*200*/IS*605* family subgroup IS*1341* (VNG_0013C, VNG_0044H, and VNG_2652H) and one protein encoded by the multi-copy IS*H2* element (VNG_0210H), belonging to the IS*H8* family (see Table S3 for IS information). All mobilome proteins, except for one (VNG_0051a), were present in the SWATH-MS assay library and none were predicted to be membrane-associated. Moreover, all produced at least one suitable tryptic peptide (≥ 7 and ≤ 30 amino acids) when digested *in silico* (Rapid Peptides Generator) (61). Notwithstanding their low CAI (Figure 5D), the high mRNA abundance (Figure 5E), and presence of TPS suggests that the mobilome proteins were not detected by virtue of being expressed at low abundance, and possibly due to post-transcriptional repression of translation by SmAP1 and asRNAs (Figure 5A). For instance, the translational repression of VNG_0112H (IS*H3* family) would be consistent with the observed pile-up of Ribo-Seq reads at the 5’ end of the transcript, which is co-located with SmAP1 binding sites and a TPS (Figure S7). Together, these observations suggest SmAP1 binding might lead to a potentially stalled ribosome-transcript complex, which may then be targeted by an endonuclease in a well-known mechanism called “No-Go” decay, as previously hypothesized for similar observations (47). The evidence provided by the atlas offered confidence for further wet lab experimental exploration. Therefore, we investigated the role of SmAP1 in regulation of IS element-mediated genome reorganization by performing long-read DNA sequencing (DNA-Seq) to quantify transposition events of each IS family in Δ*ura3*Δ*smap1* strain and its parent Δ*ura3* (Figure 6; Figure S8; Table S4; File S8). In so doing, we discovered that knocking out SmAP1 significantly decreased the overall number of transposition events (Figure 6A), and in particular transposition of the IS*4* and IS*H3* families (Figure 6B-C).

**Figure 5.**
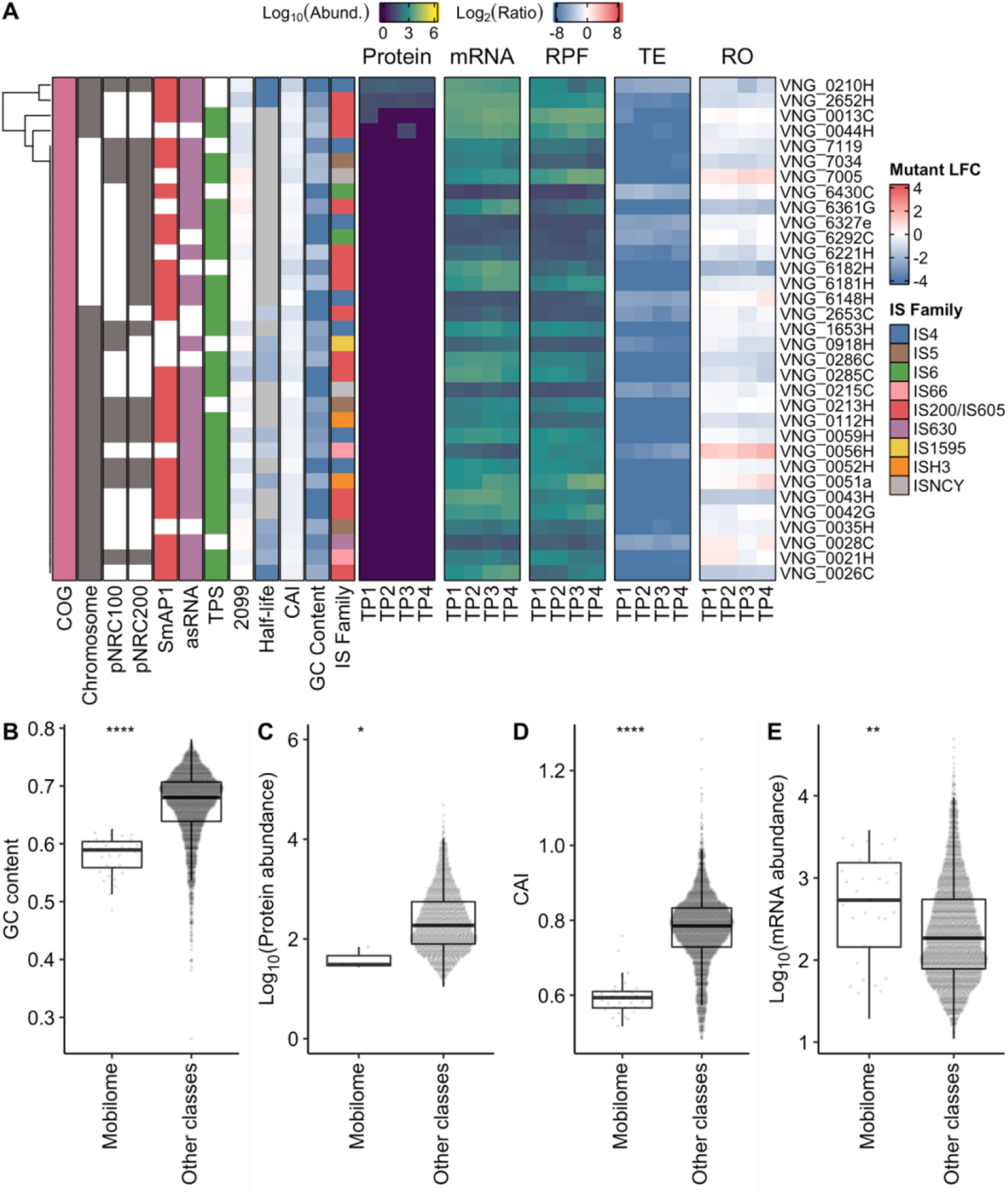
Protein and mRNA levels of mobile genetic elements. **A**. Log_10_-transformed expression profile of proteins (a pseudocount was imputed for missing values), mRNAs (TPM+1), and ribosome-protected mRNA fragments (RPF; TPM+1) with miscellaneous properties of genes classified by clusters of orthologous genes (COG) within the “Mobilome: prophages, transposons” category (pink). TE: translational efficiency; RO: ribosome occupancy; asRNAs: antisense RNA; TPS: transcript processing site; 2099: log_2_ fold change (LFC) of transcripts in the absence of RNase_2099C; TP1: early exponential growth phase; TP2: mid-exponential growth phase; TP3: late exponential growth phase; TP4: stationary phase. Box plots aid the comparison between features of genes within the “Mobilome: prophages, transposons” versus the pool of the other categories: **B**. GC content; **C**. Log_10_-transformed average protein abundance across all time points (missing values excluded); **D**. Codon adaptation index (CAI). **E**. Log_10_-transformed average mRNA levels (TPM+1) across all time points. We compared medians using the Mann–Whitney U test. * *p*-value ≤ 5×10^−2^; ** *p*-value ≤ 10^−2^; **** *p*-value ≤ 10^−4^.

**Figure 6.**
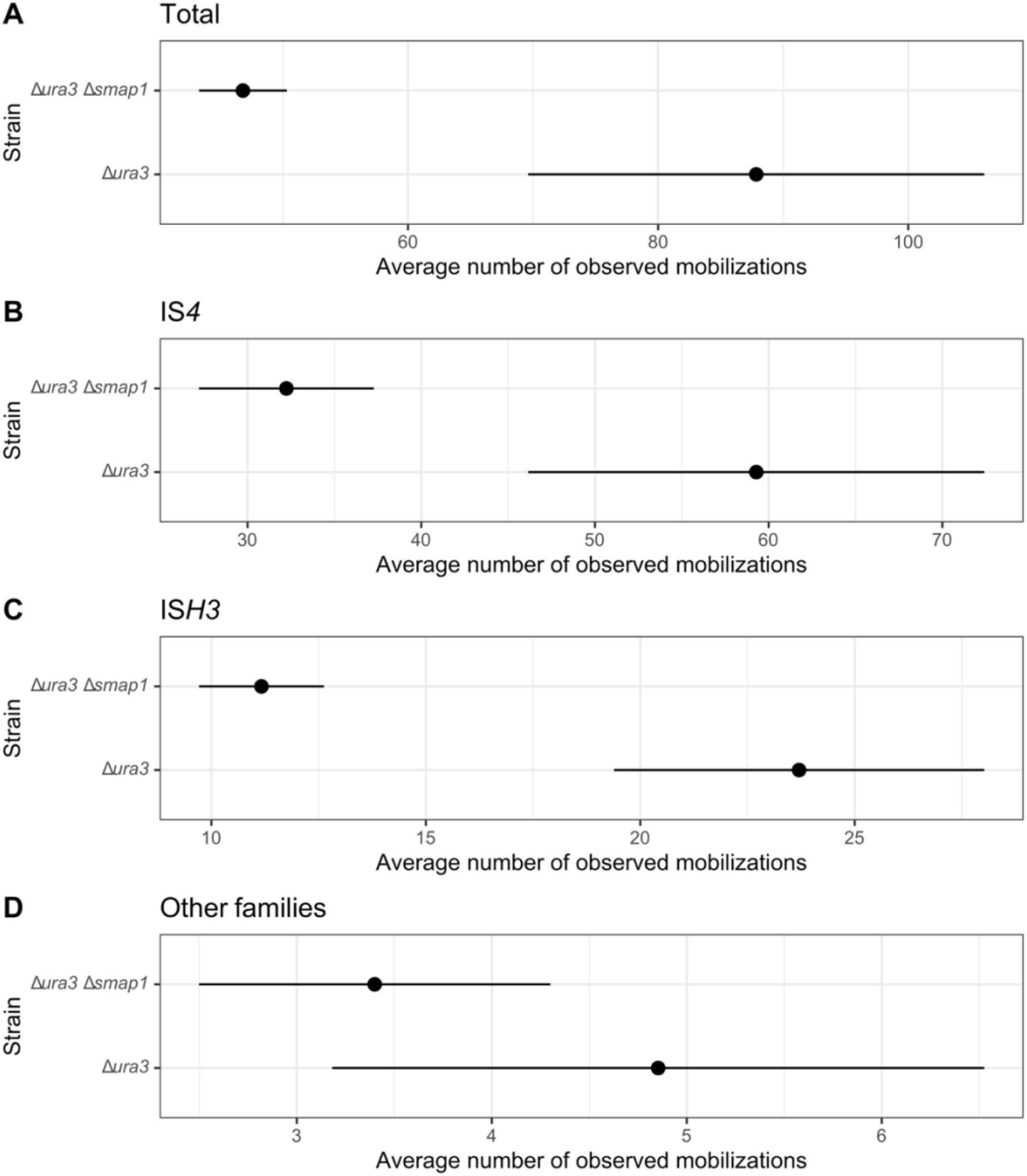
Detected mobilizations for decomposed insertion sequence families. The figure shows the average normalized number of clusters for each strain. The panels, from top to bottom, show the results for the (**A**) pool of all insertion sequences, (**B**) IS*4* family only, (**C**) IS*H3* family only, and (**D**) the other families. Black lines indicate the range of the 68% confidence interval.

#### The role of post-transcriptional regulation in governing environmental responsiveness and timing of gas vesicle biogenesis

Gas vesicles are intracellular proteinaceous organelles filled with ambient gas that may be used as buoyancy devices by halophilic archaeal cells to float to the surface to access oxygen, which has poor solubility in hypersaline water (62). The gas vesicles also act in conjunction with sensory rhodopsin-mediated phototaxis to support phototrophic energy transduction by bacteriorhodopsin (63). Hence, the biogenesis of gas vesicles is highly responsive to environmental stimuli, and in particular oxygen availability (64). Gas vesicles are made up of two structural proteins: GvpA, a monomer, and GvpC, which wraps around and stabilizes the vesicle assembled from the GvpA polymer (65). Many other proteins (GvpF-M) are involved in nucleation and biogenesis of the gas vesicle (66), a process that is regulated by GvpD and GvpE (40). The bidimensional trajectories of changes in mRNA and protein levels revealed that while the transcript levels of all *gvp* genes, including the structural proteins, increased across the four growth phases, the corresponding protein levels did not increase until the cells transitioned from mid-exponential growth phase into the stationary phase (Figure 7A), which is consistent with the timing of gas vesicle production (67). Together, the multiple levels of evidence in the *H. salinarum* NRC-1 Atlas (Figure 7B; Figure S9) supports a model (Figure 7C) that explains how the interplay of negative and positive regulation at the transcriptional, post-transcriptional, and translational levels governs the timing and environmental responsiveness of gas vesicle biogenesis.

**Figure 7.**
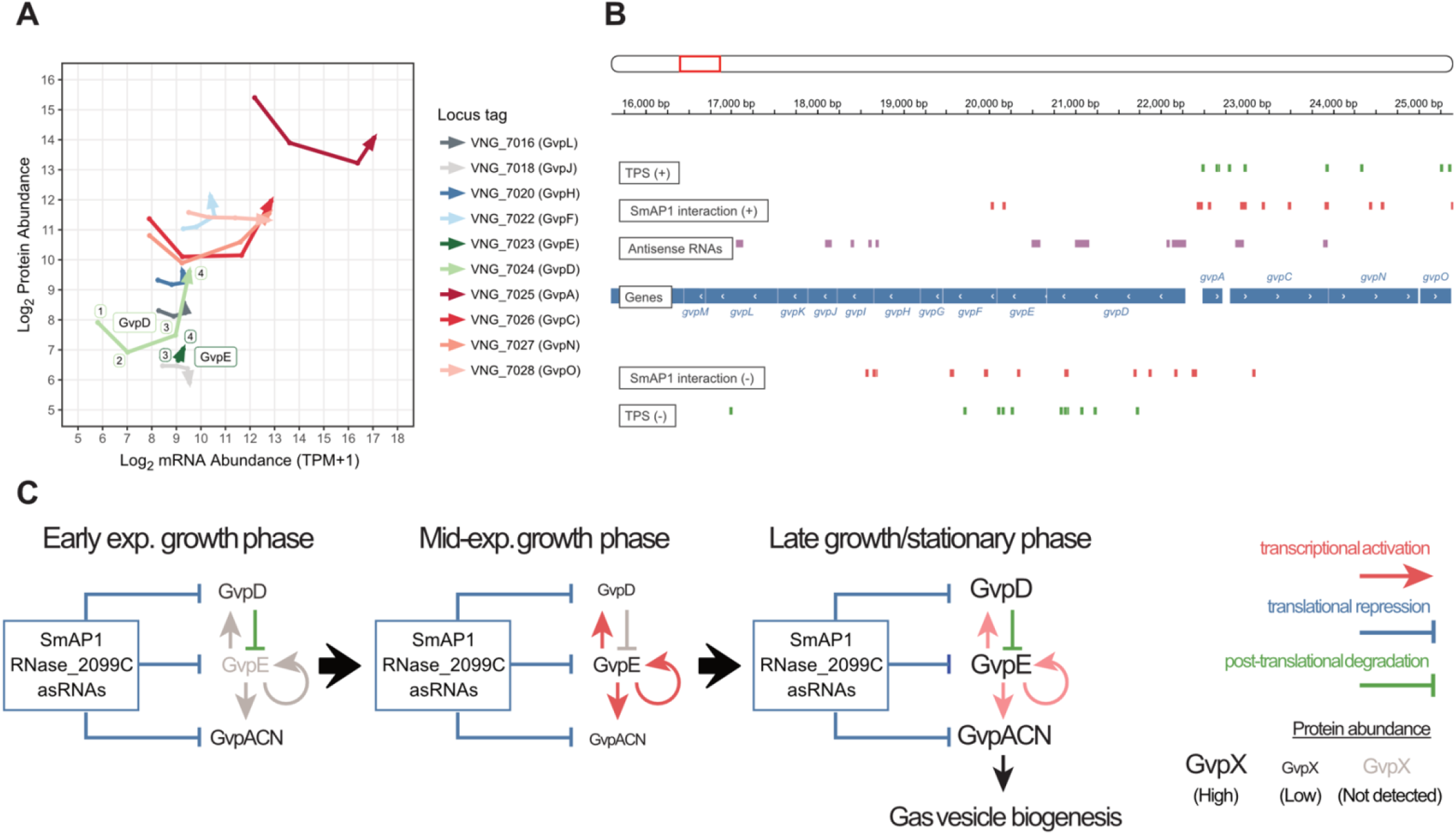
Post-transcriptional regulation of *gvp* operons. **A**. Arrows represent how each one of the gas vesicle operon genes (color-coded; protein names in parentheses) behaves regarding its log_2_-transformed protein abundance (*y*-axis) and mRNA abundance (*x*-axis) across consecutive physiological states (TP1: early exponential growth phase; TP2: mid-exponential growth phase; TP3: late exponential growth phase; TP4: stationary phase). We represent *gvpMLKJIHGFED* and *gvpACNO* operons, except for a few elements (*gvpG, gvpI, gvpK*, and *gvpM*), whose protein levels were not detected by our SWATH-MS approach. **B**. The genome browser snapshot reveals the region of *gvpDEFGHIJKLM* (reverse strand) and *gvpACNO* (forward strand) (NC_001869.1:16,000-25,500). We depict genes as blue rectangles. Tracks show various features described on the left-hand side of the panel. Green ticks represent transcript processing sites (TPS); red rectangles represent SmAP1 binding sites; purple rectangles represent annotated antisense RNAs. **C**. Time point-wise regulatory scheme of gas vesicles proteins encoded by the *gvp* cluster. Blue bars represent translational repression, red arrows represent transcriptional activation, and green bars represent post-translational degradation. Protein abundance is depicted by the font size of gas vesicle proteins (GvpX).

Based on the absolute abundance and relative change in mRNA and protein levels, we posit that *gvp* genes were constitutively transcribed across all phases of growth. But translation of *gvp* transcripts required further transcriptional activation by GvpE (68), which was prevented in early and mid-exponential growth phase by GvpD. Specifically, in the early growth phase GvpD was high in abundance and above a threshold at which it drives the degradation of GvpE (69, 70) (Figure 7AC). As cells transitioned from early to mid-growth phase, SmAP1, RNase_2099C, and asRNAs acted in concert to repress translation of *gvp* transcripts, which was especially evident in the pile-up of ribosomal footprints in the 5’ segment of the *gvpA* transcript. This putative post-transcriptional repression of translation resulted in growth-associated dilution of Gvp protein abundance, despite a steady increase at the mRNA level (Figure 7AC; Figure S10). As a consequence, GvpD protein abundance dropped below the abovementioned threshold, disrupting its ability to drive continued degradation of GvpE. This is consistent with the observation that GvpE protein was only detected in later stages of growth after GvpD abundance had decreased (Figure 7AC). Moreover, the appearance and subsequent increase in abundance of GvpE post-mid-exponential growth phase likely resulted in transcriptional activation of all *gvp* genes (Figure 7AC). Indeed, mRNA levels of all *gvp* genes increased by >4-fold in mid-exponential growth phase (despite active cell division), unlike the moderate (≈2-fold) albeit steady increase observed in early and late phases of growth (Figure 7A). The transcriptional activation of all *gvp* genes likely overcame SmAP1, RNase_2099C, and asRNA-mediated post-transcriptional repression to upregulate translation via increased ribosomal read through (Figure 7C; Figure S10). The resulting dramatic increase in abundance of proteins GvpN and GvpO, as well as the chaperone GvpF, potentially triggered the recruitment of GvpA to initiate gas vesicle assembly (66). Concomitantly, in the stationary phase, GvpD protein level increased above the threshold, likely restoring GvpE degradation, thereby disrupting transcriptional activation of *gvp* genes, and potentially terminating further translation of gas vesicle proteins (Figure 7C). So, in essence, the interplay between GvpD-mediated degradation of GvpE, transcriptional activation of *gvp* genes by GvpE, and post-transcriptional repression of translation of *gvp* genes (likely mediated by SmAP1, asRNAs, and RNase_2099C), together modulated timing of gas vesicle biogenesis. In this scheme, subtle changes in interplay across the different levels of regulation could drive rapid initiation or termination of gas vesicle biogenesis, given that the transcripts and the monomeric structural proteins are maintained at relatively high abundance, but the regulatory (GvpD and E) and some accessory proteins (e.g., GvpJ and L) are at significantly lower abundance across all growth phases.

## DISCUSSION

This study has uncovered that a strikingly large proportion of protein-coding genes (54%) in the *H. salinarum* NRC-1 genome are potentially post-transcriptionally regulated. Notably, this estimate of the scale of post-transcriptional regulation is based on compilation of evidence from a limited set of contexts (i.e., primarily standard growth conditions). It is noteworthy that comparison of absolute and relative abundance changes in mRNA and protein levels just over batch culture growth has provided evidence for post-transcriptional control of 7% of all protein-coding genes. Different sets of genes were previously reported to have discordant relationship between mRNA and protein levels in other environmental contexts such as shifts in oxygen tension (44) and exposure to gamma irradiation (45). In response do gamma irradiation, 47 upregulated transcripts had direction of change incompatible with their respective proteins. Of those, only five are included in the set of 188 putative post-transcriptionally regulated genes identified by the present study. Together, these observations illustrate the importance of environmental context in characterizing genome-wide implications of post-transcriptional regulation. Similarly, we have surveyed just three mechanisms (SmAP1, asRNAs, and one RNase) that provide likely mechanistic explanation for post-transcriptional regulation of 430 out of 966 transcripts (45%) with TPS. This suggests that the remaining TPS-associated 536 transcripts are potentially post-transcriptionally regulated by other mechanisms, including endoribonucleases, trans-acting antisense RNAs and small regulatory RNAs (sRNAs) that were not surveyed in this study. Although, prior work has suggested a limited role for trans-acting antisense RNAs and sRNAs in archaeal regulation (71). Nonetheless, we can expect many more genes in the *H. salinarum* NRC-1 genome to be subject to post-transcriptional regulation, especially in ecological contexts that require rapid physiological state transitions for environmental acclimation.

Transcriptome-wide binding analysis with RIP-Seq implicated a global role for SmAP1 in post-transcriptional regulation of at least 397 genes. Action of SmAP1 in *H. salinarum* NRC-1 appears to have mechanistic similarity to its counterparts in other archaea and also to Hfq in bacteria, such as preferentially targeting AU-rich sequences, and regulating itself (35). Autoregulation by the bacterial ortholog of SmAP1, Hfq, has also been reported previously in *E. coli* (72, 73) and *Sinorhizobium meliloti* (74). By reviewing RIP-Seq results from studies in other archaea we discovered that SmAP1 also binds to its own transcript in *S. solfataricus* (SSO6454) (33). The absence of evidence for autoregulation of SmAP1 in *H. volcanii* (31) is likely a technical artefact because the microarray used for RIP-ChIP interrogated binding to only non-coding RNAs, and did not include probes for coding genes, including the SmAP1 CDS (HVO_2723). Further, the genes targeted by SmAP1 also bear functional similarity with other organisms wherein SmAP1 has been implicated in the regulation of motility (32, 75) and its ortholog has been implicated in regulation of transposition (59, 60). Notably, of the 32 non-redundant IS element-encoded proteins (Table S3) with above-average mRNA levels, only four were detected by SWATH-MS in this study, suggesting they were all post-transcriptionally repressed. By analyzing proteomics data from PeptideAtlas (76, 77) and PRIDE (78), including PXD003667 (79) and PXD015192 (80), we confirmed that 50% of the 32 transposases have been previously detected, depending on techniques and biological conditions. In addition, except for VNG_0051a, we established that these proteins bear the features required for detection by SWATH-MS. With that reasoning, we posit that the lack of detection of transposases in this study is due to their low abundance or complete absence. Together these findings make a compelling case that translation of IS element-encoded transposases, and therefore transposition of mobile genetic elements, is post-transcriptionally regulated. Translational inhibition of transposases might have evolved as a fail-safe measure to prevent transposition in most contexts and allow their rapid activation in stressful environmental contexts, wherein benefits of genome reorganization could outweigh their deleterious effects (81).

Notwithstanding their mechanistic and functional similarities with counterparts in other archaea and even bacteria, we discovered that consequences of SmAP1-mediated regulation of transposition by some families of IS elements in *H. salinarum* NRC-1 are counterintuitive. Specifically, while we had expected that SmAP1 may likely repress translation of transposase transcripts, to our surprise we discovered that deletion of SmAP1 resulted in decreased frequency of transposition by IS elements of the IS*4* and IS*H3* families, which brought to fore two outstanding questions. First, in addition to directing targeted post-transcriptional processing and repression of transcripts, (how) does SmAP1 also mediate transposition by IS elements? And second, despite targeting AU-rich sequences how do SmAP1 and its counterparts accomplish regulation of specific subsets of target genes in a context-specific manner? While the first question will need further investigations into the mechanisms of SmAP1 action on transposition events, our integrated analysis has provided some clues to address the second question, such as evidence that SmAP1 might act in concert with other post-transcriptional regulatory mechanisms, viz., asRNAs and RNase_2099C to gain specificity for transcripts. So while SmAP1 appears to be expressed constitutively and maintained at median abundance (Figure S2B), its mode and target of action may be governed by other factors, such as conditional expression of asRNAs, which could possibly guide SmAP1 action on specific transcripts in a similar manner to its bacterial counterpart (24). Indeed, in *H. volcanii* the global oxidative stress response upregulates asRNAs with consequential downregulation of specific transposase mRNAs, especially of the IS*4* family (71). For example, SmAP1 and an asRNA may jointly regulate transposition events by binding to the 5’ end of TnpB (VNG_0042G) transcript to repress translation of this putative RNA-guided endonuclease, which is encoded by IS*H39* (IS*200*/IS*605* family) and possibly part of the transposition apparatus (Figure S11) (82, 83). Thus, SmAP1-mediated post-transcriptional regulation of mobile elements appears to have pleiotropic consequences depending on the IS family, with a repressive role for IS*200*/IS*605*, as reported previously for *S. enterica* (60), and an enhancer role for IS*H3* and IS*4*. Indeed, SmAP1 might facilitate translation of transcripts, considering its hairpin-melting potential (84) and its role as a recruiter for translational complex subunits (85).

The current study has revealed extensive interplay of post-transcriptional regulation with regulation at other levels of information processing, which may mediate rapid adaptive responses to environmental change (e.g., genome reorganization by triggering transposition of IS elements, and vertical relocation by activating gas vesicle biogenesis). In the case of gas vesicle biogenesis, we observed that the high abundance and relative increase in transcript levels of the gas vesicle structural genes did not manifest in increased protein levels until the post-transcriptional repression of translation was overcome in later stages of growth, which is associated with stressful conditions including anoxia and nutrient limitation. Previously, we had demonstrated that RNase_2099C is transcriptionally co-regulated with genes of the aerobic physiologic state but acts on transcripts of the anaerobic state (21). In this arrangement, the interplay of RNase_2099C with transcriptional regulation generates an efficient state transition switch. For instance, RNase_2099C-mediated repression of positive transcriptional autoregulation (RPAR) enables rapid shutdown of ATP-consuming K^+^ uptake to conserve energy under anoxic conditions with high potassium availability. Gas vesicle biogenesis (response to light and oxygen) appears to be regulated in a similar set up albeit with an expanded set of players. Specifically, the interplay of GvpD-mediated degradation of GvpE, GvpE-mediated transcriptional activation of *gvp* genes, and post-transcriptional repression of gas vesicle protein synthesis through potential interplay of SmAP1, RNase_2099C, and asRNAs is likely critical for mediating rapid initiation and termination of gas vesicle biogenesis. The genome-wide atlas reveals that a large proportion of genes in the *H. salinarum* NRC-1 genome is likely subject to such post-transcriptional regulation, and as such it will serve as an interactive hypothesis generator to drive in-depth characterization of specific mechanisms of rapid environmental acclimation.

## MATERIALS AND METHODS

### Strains, media, and growth conditions

We grew *Halobacterium salinarum* NRC-1 in complex media (CM; 250 g/L NaCl, 20 g/L MgSO_4_•7H_2_O, 3 g/L sodium citrate, 2 g/L KCl, and 10 g/L bacteriological peptone). Mutant strains, Δ*ura3* and Δ*ura3Δsmap1*, had their media supplemented with uracil (50 µg/mL). Vector harboring strains, wtp-pMTF-cMyc and wtp-pMTF-SmAP1-cMyc, had their media supplemented with mevinolin (20 µg/mL). All the cultures were grown at 37 ºC, under light, and with constant agitation of 125 RPM (otherwise specified). For cloning steps, we used *Escherichia coli* DH5α grown in lysogeny broth (LB; 10 g/L tryptone, 5 g/L yeast extract, 10 g/L NaCl, pH 7.5) at 37 ºC and under constant agitation. Carbenicillin (50 µg/mL) was added to LB when necessary.

### Construction of SmAP1 knockout strain and SmAP1 tagged strain

SmAP1 knockout strain (Δ*ura3*Δ*smap1*; Δ*VNG_1673G*Δ*VNG_1496G*) was constructed from a parent Δ*ura3* strain (Δ*VNG_1673G*) by using the pop-in/pop-out method with two-step selection by mevinolin and 5-fluoroorotic acid (5-FOA) (86). Polymerase chain reaction (PCR) was used to confirm the genotype of null mutants selected by 5-FOA (Table S5). We evaluated the growth curve phenotype (Figure S12) by culturing strains in CM supplemented with uracil (50 μg/ml) at 37 °C and 125 RPM.

To create the recombinant protein SmAP1-cMyc, we used the pMTF-cMyc vector (4). The SmAP1 encoding gene (VNG_1496G) was amplified (Table S5) and purified using QIAquick PCR Purification (QIAGEN). The amplification product was cloned into the vector pMTF-cMyc, upstream to the region encoding 13-cMyc tag. The procedure was carried out by digesting pMTF-cMyc with endonucleases NdeI and BamHI (Fermentas) with further ligation of *smap1* amplicon by T4 DNA ligase (Fermentas). The clone was transformed into *E. coli* DH5α and confirmed by PCR and Sanger sequencing. Vectors were extracted and transformed into *H. salinarum* NRC-1 strain to create strains wtp-pMTF-SmAP1-cMyc (SmAP1-cMyc overexpression) and wtp-pMTF-cMyc (cMyc-overexpression).

### SmAP1-RNA co-immunoprecipitation

*H. salinarum* strains wtp-pMTF-SmAP1-cMyc and wtp-pMTF-cMyc were grown until they reached OD_600nm_ ≈ 0.75. We centrifuged 20 mL of cell culture at 3,700 RCF for 10 minutes and resuspended cells in 12 mL of basal solution (CM without bacteriological peptone). The cellular suspension solution was transferred to Petri dishes, on ice, and submitted to 800×100 µJ/cm^2^ ultraviolet (UV) radiation inside a UVC 500 Crosslinker (Amersham Biosciences). It was carefully transferred to 50 mL tubes and centrifuged at 3,700 RCF for 15 minutes at 4 ºC. Cells were resuspended in 1 mL of lysis solution (1x PBS, 0.1% SDS, 0.5% deoxycholate, 0.5% NP-40, proteinase inhibitor—1 tablet for 100 mL (Sigma S8830), RNaseOUT inhibitor—2 µL/10 mL (Invitrogen)) and ice incubated for five minutes. The suspension was centrifuged at 10,000 RCF for five minutes at 4 ºC. The supernatant was separated and incubated with 10 µL of Dynabeads M-450 anti-mouse IgG (Invitrogen #11041) for 10 minutes, at 4 ºC, to remove spurious interactions. After incubation, the solution was centrifuged at 10,000 RCF for five minutes at 4 ºC. The supernatant was incubated overnight, under constant agitation, at 4 ºC, with 60 µL of anti-cMyc (antibody) coated beads (Sigma M4439). Beads were immobilized using a magnetic rack and washed twice using 1 mL of lysis solution, followed by two rounds of washing with 1 mL of saline solution (5x PBS, 0.1% SDS, 0.5% deoxycholate, 0.5% NP-40), and finally washed with 1 mL of Tris-EDTA (TE buffer). Beads were resuspended in 100 µL of TE and incubated at 65 ºC for 10 minutes. The suspension was centrifuged at 14,000 RCF for 30 minutes at 25 ºC. We added 120 µL of TE/SDS (SDS 0.1%) to the supernatant and incubated it for 30 minutes at 65 ºC. Two aliquots were separated: i) one destined to the western blot assay; and ii) another destined to the RNA isolation prior to sequencing.

### SmAP1-cMyc western blot assay

We verified the presence of the SmAP1 protein in the co-immunoprecipitated samples using the western blot assay. Aliquots were added of sample buffer (30% glycerol (v/v), 9.2% SDS (w/v), 1% bromophenol blue (w/v), 20% β-mercaptoethanol (v/v), 0.25 M Tris-HCl pH 7.0) and denatured at 95 ºC for five minutes. Denatured samples (20 µL) were submitted to 10% polyacrylamide gel electrophoresis (SDS-PAGE). PageRuler Prestained Protein Ladder (Fermentas) was used as weight marker and transference control. Gel and Hybond ECL nitrocellulose membrane (GE) were dipped in transfer buffer for 10 minutes.

The membrane transfer was performed at 100 V for one hour. The membrane was washed with PBS-T (0.1% Tween 20 (v/v)) and incubated in PBS-T with milk at room temperature for one hour. After the blocking step, the membrane was quickly washed twice with PBS-T. The primary antibody (anti-cMyc) was diluted (1:3,000) in PBS-T, and incubation was carried out at 4 ºC, under constant agitation, overnight. The membrane was rewashed with PBS-T and incubated in PBS-T at room temperature, under constant agitation for 15 minutes. The secondary antibody (anti-mouse IgG-peroxidase - Sigma A4416) was diluted (1:3,000) in PBS-T, and incubation was carried out at room temperature, under constant agitation, for one hour. The membrane was quickly washed twice using PBS-T and incubated in PBS-T at room temperature, under constant agitation, for 15 minutes. We used the reagents ECL Western Blotting Detection (GE) to develop the membrane, and images were obtained using ChemiDoc XRS+ (Bio-Rad).

### SmAP1 RIP-Seq and data analysis

The co-immunoprecipitated RNA samples were submitted to protein digestion using proteinase K (Fermentas) and purified using the MinElute Reaction Cleanup Kit (QIAGEN) with a DNase treatment step. We quantified the RNA samples using Quant-iT RiboGreen RNA Assay (Invitrogen) and prepared them for sequencing using the TruSeq mRNA Stranded kit (Illumina). Before sequencing, to equalize the concentrations, quantification was performed by using the KAPA Library Quant kit (Kapa Biosystems). Samples were sequenced using the MiSeq Reagent v2 kit (Illumina) for 50 cycles, using the single-end mode, in a MiSeq instrument (Illumina).

We processed the sequenced libraries using the ripper pipeline (Table S6) to obtain putative SmAP1 binding regions. Briefly, the software: i) trims the bad quality ends and adapters from reads using Trimmomatic (87); ii) aligns trimmed reads to the reference genome (NCBI Assembly ASM680v1) using HISAT2 (88) without gaps, splicing, or soft-clipping; iii) converts alignment files from SAM to BAM format using SAMtools (89); iv) adjusts multi-mapping reads using MMR (90); v) computes single-nucleotide resolution transcriptome signal using BEDtools (91); vi) computes a coordinate-wise log_2_ fold change between co-immunoprecipitated samples relative to control samples and identify regions with at least ten consecutive nucleotides satisfying log_2_ fold change ≥ 1. Interaction regions for two biological replicates (BR1 and BR2) were merged, since their intersection of SmAP1-bound genes had a 3.8-fold enrichment over the expected value (observed: 157; expected: 41.44; *p*-value = 3.14×10^−71^). We tested the fold enrichment significance by using the SuperExactTest::MSET function (92).

### Preparation and acquisition of proteomics samples

Sample preparation and data acquisition for the time-course measurements of the *H. salinarum* proteome were performed as described in Kusebauch et al. (in preparation). *H. salinarum* NRC-1 was cultured in CM. Cultures were grown in triplicate (37°C, shaking at 220 RPM) and illuminated (≈20 μmol/m^2^/sec) in Innova 9400 incubators (New Brunswick). Cultures were harvested at four time points: early exponential phase (OD_600nm_ = 0.2; 14.3 hours), mid-exponential phase (OD_600nm_ = 0.5; 21.5 hours), late exponential phase (OD_600nm_ = 0.8; 28.8 hours), and stationary phase (40.8 hours). Cells were collected by centrifugation (8,000 x g, 2 minutes, 4°C). Cell pellets were resuspended in Milli-Q water and disrupted at 4°C using ceramic beads (Mo Bio Laboratories) and a Precellys 24 homogenizer (Bertin Corp). Protein content was determined by bicinchoninic acid assay (BCA) (Thermo-Fisher). Proteins were reduced (5mM Dithiothreitol (DDT, 45 minutes, 37 °C)), alkylated (14 mM iodoacetamide (30 minutes, room temperature, darkness)), and digested with trypsin (1:50 enzyme:substrate ratio, 37°C, 16 h). Samples were desalted with tC18 SepPak cartridges (Waters). Sample analysis was performed on a TripleTOF® 5600+ system equipped with a Nanospray-III® Source (Sciex) and an Eksigent Ekspert™ nanoLC 425 with cHiPLC® system in trap-elute mode (Sciex). Peptides were separated with a gradient from 3% to 33% of 0.1% formic acid in acetonitrile (v/v) for 120 minutes. Data were collected in MS/MS^ALL^ SWATH™ acquisition mode using 100 variable acquisition windows.

### SWATH-MS data analysis

SWATH-MS data were analyzed with the Spectronaut software (version 15.5.211111.50606) and an assay library for *H. salinarum* NRC-1 reported in Kusebauch et al. (in preparation). SWATH .Wiff raw data files were converted to HTRMS files with the Spectronaut HTRMS converter (15.5.211111.50606). Data extraction mass tolerance (MS1 and MS2) was set to dynamic with a correction factor of 1. Dynamic extracted ion chromatogram (XIC) RT window was enabled with a correction factor of 1 and local (non-linear) RT regression. Decoy assays were dynamically generated using the scrambled decoy method and library size fraction set to 1. The identification was performed using the normal distribution estimator with precursor identification results with *q*-value (false discovery rate; FDR) < 0.1 and protein identification results with a *q*-value (FDR) < 0.01. Quantification was performed with interference correction enabled, MS2 ion peak areas of quantified peptides were summed to estimate protein peak areas, and area as quantity type selected. Identified precursor quantities were normalized using the Spectronaut built-in global normalization function (median). The four time points in this study were defined as four conditions in the condition setup. We used Spectronaut’s protein quantification and proDA (93) to perform differential expression analysis of proteins. We computed the contrasts of interest and set up |log_2_ fold change| ≥ 1 and adjusted *p*-value < 0.05 as the criteria to determine differentially expressed proteins.

### Non-redundant reference transcriptome

Many annotation efforts for *H. salinarum* NRC-1 have been made available since the publication of its genome assembly (49). Consequently, cross-referencing findings from publications using different sources has become a challenging and time-consuming task. Moreover, the genome presents redundancy in terms of (quasi)identical paralogs, most of them found within plasmid repetitive regions (94) and contained within multi-copy insertion sequences (95). To solve the problem of the annotation multiplicity and gene redundancy, we extracted coding and non-coding sequences (tRNAs, rRNAs, Signal Recognition Particle RNA, and RNase P) from different annotation sources for *H. salinarum* NRC-1 and R1 strains (Table S1) and clustered them using CD-HIT (96). Coding and non-coding genes with at least 95% and 99% global amino acid and nucleotide identity, respectively, were grouped and represented by a single entity anchored by the sequence and locus tag given by the latest large-scale annotation effort for *H. salinarum* NRC-1 (50). We only considered sequences represented in this annotation. We also collected and parsed clusters of orthologous genes (COG) (97) to functionally categorize the non-redundant reference transcriptome, and classified insertion sequence families using ISfinder (56) and ISsaga (98) platforms. The code to reproduce this annotation simplification effort is available on GitHub (see halo_nr_tx in Table S6).

### Transcriptome analysis

We retrieved RNA-Seq and Ribo-Seq data from a *H. salinarum*’s growth curve experiment available at NCBI SRA under accession PRJNA413990 (42). The samples are the same for which the proteome data was generated, as explained previously. We quantified all the RNA-Seq libraries by mapping them against the *H. salinarum* NRC-1 non-redundant reference transcriptome using kallisto (99) facilitated by the use of the pipeline runKallisto (Table S6). We performed differential expression analysis for the RNA-Seq and Ribo-Seq time course experiment (42) using DESeq2 (100). Only genes satisfying |log_2_ fold change| ≥ 1 and adjusted *p*-value < 0.05 were considered differentially expressed. We generated the transcriptome coverage signal for genome browsing using the frtc pipeline (101) (Table S6). Briefly, the tool trims reads using Trimmomatic (87); aligns them to the reference genome (NCBI Assembly ASM680v1) using HISAT2 without splicing (88); adjusts multi-mapping instances using MMR (90); and computes the genome-wide coverage using deepTools2 (102).

We performed differential expression analysis of strain Δ*RNase_2099C* by reanalyzing data from (21), deposited in Gene Expression Omnibus (GEO) under accession GSE45988. Briefly, we used limma (103) to process the data and computed the Δ*RNase_2099C* vs. *Δura3* contrast controlling for the growth curve time point effect. We only used mid-exponential (OD_600nm_ ≈ 0.4) and late-exponential (OD_600nm_ ≈ 0.8) growth phase data. Only genes satisfying |log_2_ fold change| ≥ 1 and *p*-value < 0.05 were considered differentially expressed.

### Inference of putative post-transcriptionally regulated genes

We relied on transcriptome and proteome quantitation to infer putative post-transcriptionally regulated genes. For that, we developed two methods: i) the absolute abundance-based approach, in which we identified genes producing simultaneously high mRNA levels (transcripts per million, TPM, in the upper quintile) and low protein abundance (lower quintile or undetected); and ii) the relative abundance-based approach, in which we inspected differentially expressed genes in physiological state transitions having mRNA levels being upregulated whilst protein levels being downregulated.

We further inspected genes identified by the absolute abundance-based approach, whose proteins were not detected, to remove entries likely missed due to technical limitations. After manual inspection, we removed potential transmembrane proteins (as these are difficult to be detected), proteins not represented in the assay library due to the lack of suitable peptides for detection by SWATH-MS (e.g., hydrophobicity, peptide length), and proteins not represented in the assay library due to differences in annotation versions. To be considered a transmembrane protein, we first conducted a transmembrane domain prediction for all the entries encoded by the non-redundant transcriptome using TOPCONS webserver (104). We manually inspected the results and evaluated the “consensus prediction probability” of transmembrane regions. We required proteins to have at least one transmembrane domain with a considerable extension satisfying probability ≥ 0.9. To aid our judgement, we also pondered empirical evidence (105, 106) and functional annotation. This approach identified 117 genes with expressive mRNA and undetected proteins with a high likelihood of being post-transcriptionally regulated (File S3).

### Long-read DNA sequencing and analysis

*H. salinarum* strains Δ*ura3* and Δ*ura3*Δ*smap1* were grown in CM supplemented with uracil until OD_600nm_ ≈ 0.5. Aliquots of 2 mL of cell cultures were submitted to DNA extraction using DNeasy Blood & Tissue kit (QIAGEN). DNA samples were quality checked and genotyped using PCR to confirm strains (Table S5). We prepared the samples for long-read DNA sequencing using the MinION platform (Oxford Nanopore Technologies, ONT). Libraries were prepared using SQK-LSK108 (ONT) combined with EXP-NBD103 (ONT) to allow multiplexing. The experiment was run using MinION Mk1B (ONT) in a FLO-MIN106 (ONT) flow cell for 48 hours. Raw data were demultiplexed using Deepbinner (107), and base called by Guppy (ONT). Quality checking was done using Filtlong (Table S6), and adapter trimming was performed using Porechop (Table S6).

We used NGMLR (108) to align reads to a modified version of reference genome, which excludes long duplications (NC_002607.1:1-2,014,239, NC_001869.1:1-150,252, NC_002608.1:112,796-332,792). To identify structural variations (SV), the alignments were processed with Sniffles (108), and the VCF files were filtered to keep only insertions and deletions. The sequences of detected SVs were compared to *H. salinarum* NRC-1 annotated insertion sequences using BLAST (109). Insertions and excisions were only annotated if satisfying the threshold of at least 75% identity, 80% coverage considering both query and subject. These criteria were based on the 80-80-80 rule proposed by (110), but slightly loosened because of Nanopore intrinsic high error rates.

We applied a clustering approach for neighbor elements to avoid overestimating the number of identified SVs. SVs of the same class (insertion or excision), caused by the same element, and starting within 50 base pairs of distance from each other, were combined into a single cluster having a mean start point and a support index based on the number of occurrences. Dividing this number of occurrences (*e*) by the local read coverage (25-nucleotide bidirectional flank) (*c*) allowed us to classify SV clusters in three categories: i) When *e/c* ≤ 0.1, the cluster is defined as relatively rare in the population; ii) When 0.1 < *e/c* ≤ 0.5, it is common; iii) When *e/c* > 0.5, it is characterized as predominant, indicating this SV might be fixed in the population genomes.

We computed the total number of clusters of insertions and excisions for each of the libraries and added them up before normalizing the values based on each sample’s total of aligned reads. To normalize, we identified the library with the biggest number of aligned reads and adjusted the others to be comparable. The mean value for normalized counts was computed for both Δ*ura3*Δ*smap1 and* Δ*ura3* and compared using a confidence interval of 68% (see Table S6 for code).

### Enrichment analysis and average comparison

To detect enriched features (e.g., SmAP1 binding, asRNA, and TPS) within groups of genes, we performed enrichment analysis using the hypergeometric test from R software (stats::phyper function). To compare the average of features (e.g., half-lives, CAI, GC, and Δ*RNase_2099C* log_2_ fold change (LFC)) between groups of genes, we used the nonparametric Mann–Whitney U test from R software (stats::wilcox.test function). The significance cutoff of our choice for both statistical tests was *p*-value < 0.05.

### Data collection from miscellaneous sources

We gathered and parsed data from several sources. We collected antisense RNA (asRNA) data from Table S4 of (46). We obtained transcript processing sites (TPS) from Table S1 of (47). Redundancy was removed by collapsing asRNAs and TPS of identical and (quasi)identical transcripts. We obtained half-lives from a microarray experiment (43). The redundancy was removed by computing the average half-lives of identical and (quasi)identical genes. We computed the codon adaptation index (CAI) (111) using the coRdon::CAI function (see coRdon in Table S6), taking as input the 5% most abundant proteins according to our proteomics approach. We computed the GC content (guanine-cytosine content) using the Biostrings::letterFrequency function.

### H. salinarum *NRC-1 multi-omics Atlas portal*

We developed the *H. salinarum* NRC-1 multi-omics Atlas portal by integrating existing components to new resources. Legacy data is stored in an SBEAMS MS SQL Server database which supplements the main MySQL database. A web service API implemented in Python and Flask provides uniform access to these resources. We implemented the web-based user interface using the Javascript framework Vue.js (see Table S6 for code). We built the heatmap interface with the help of InteractiveComplexHeatmap (112), ComplexHeatmap (113), and Shiny R packages. We built the genome browser by using igv.js (114). Data used to generate heatmaps were prepared as described in previous sections with an additional step for scale adjustment allowing a graphical representation of disparate multimodal omics sources. The quantile normalized data is also available along with the non-normalized data (File S1). The web portal is available at http://halodata.systemsbiology.net.

## Supporting information

Supplemental File Titles

## Data and code availability

SmAP1 RIP-Seq raw data (FASTQ format) and DNA-Seq data (demultiplexed, base called, and trimmed; FASTQ format) were deposited in NCBI’s Sequence Read Archive and are publicly available under the BioProject accession PRJNA808788. Raw DNA-Seq data (FAST5 format) is available at Zenodo under the digital object identifier 10.5281/zenodo.6303948 (accession 6303948). The code used in this study is available on GitHub in multiple repositories (see Table S6 for links and description).

## CREDIT AUTHORSHIP CONTRIBUTION STATEMENT

APRL: Methodology, Software, Validation, Formal analysis, Investigation, Data Curation, Writing — Original Draft, Writing — Review & Editing, Visualization; UK: Methodology, Investigation, Formal analysis, Writing — Review & Editing; LSZ: Methodology, Investigation; WJW: Software, Data Curation, Visualization; JPPA: Methodology, Validation, Formal analysis, Investigation, Data Curation, Writing — Review & Editing; ST: Software, Data Curation, Writing — Review & Editing, Visualization; ALGL: Conceptualization, Writing — Review & Editing, Supervision; JVGF: Conceptualization, Writing — Review & Editing, Methodology, Investigation; RZNV: Conceptualization, Validation, Writing — Review & Editing, Supervision; RLM: Conceptualization, Resources, Writing — Review & Editing, Supervision, Project administration, Funding acquisition; TK: Conceptualization, Resources, Supervision, Project administration, Funding acquisition; NSB: Conceptualization, Resources, Writing — Original Draft, Writing — Review & Editing, Visualization, Supervision, Project administration, Funding acquisition.

## ACKNOWLEDGMENTS

We thank Dr. Alessandro de Mello Varani for helping us with insertion sequence family annotation; Silvia Helena Epifânio and Min Pan for the laboratory technical support; Catarina dos Santos Gomes for helping in the execution of long-read DNA sequencing; Dr. Elisabeth Wurtmann for helping with the RIP-Seq assay standardization.

## SUPPLEMENTAL TABLES

Table S1 | Annotation sources for constructing the *Halobacterium salinarum* NRC-1 non-redundant transcriptome and a loci dictionary.

Table S2 | Comparison of Pearson correlation coefficient computed for protein and mRNA abundance throughout the growth curve. We compared the coefficients using Zou’s confidence interval method implemented in the cocor package. Subscripts *P* and *m* refer to protein and mRNA levels for indicated time points. Uppercase letters (A-F) refer to panels in Figure 2. * Δ*R* stands for the subtraction between the two coefficients (e.g., *R*_TP1(A)_ - *R*_TP2(B)_). A confidence interval (CI) of Δ*R* spanning zero is not significant. Coefficients diverge slightly from those presented in the main text due to technical differences between the comparative approach and classic correlation method implementations. TP1: early exponential growth phase; TP2: mid-exponential growth phase; TP3: late exponential growth phase; TP4: stationary phase.

Table S3 | The non-redundant set of insertion sequences in *Halobacterium salinarum* NRC-1. We obtained insertion sequence families from ISfinder and ISsaga, and the transposition mechanisms from Siguier et al. (2015).

Table S4 | Summary of the transposition detection assay. ^a^ Number of identified insertion clusters.

^b^ Number of identified excision clusters. ^c^ Number of reads aligned to the reference genome. ^d^

Sum of insertion and excision clusters normalized by the library with the highest number of aligned reads.

Table S5 | List of primers used in this study.

Table S6 | In-house and third-party GitHub repositories cited in this study.

## SUPPLEMENTAL FIGURES

Figure S1 | Quality assurance of co-immunoprecipitated samples. **A**. Western blot of samples extracted from strains expressing plasmids for cMyc and cMyc-tagged SmAP1 (see lane titles for labels). The expected molecular weight of the cMyc-tagged SmAP1 complex is 37 kDa. BR: Biological replicate. **B**. Polymerase Chain Reaction (PCR) of RNA-purified samples treated with DNase. M: Ladder; 1: Positive control (genomic DNA amplified using 19-fwd and 20-rev primers with a predicted amplicon size of 85 bp); 2-5: cMyc BR1, cMyc BR2, SmAP1-cMyc BR1, and SmAP1-cMyc BR2 (amplified using 19-fwd and 20-rev primers); 6: Positive control (genomic DNA amplified using 63-fwd and 64-rev primers with a predicted amplicon size of 450 bp). 7-10: cMyc BR1, cMyc BR2, SmAP1-cMyc BR1, and SmAP1-cMyc BR2 (amplified using 63-fwd and 64-rev primers).

Figure S2 | SmAP1 features. **A**. SmAP1 binding is conditioned to the GC content of transcripts. The reduced GC content of transcripts is a property influencing SmAP1 binding. We compared medians using the Mann–Whitney U test. **** *p*-value ≤ 10^−4^. **B**. Time course view of protein, ribosome-protected mRNA fragments (RPF; TPM+1), and mRNA levels (TPM+1). Vertical bars represent the standard error computed using at least six replicates for proteins and three replicates for mRNA and RPF. **C**. Functional categories of transcripts bound to SmAP1. The panel shows how many genes have transcripts bound to SmAP1, considering each category of COG (clusters of orthologous genes). The left-hand side panel shows categories with no more than 25 genes with SmAP1-bound transcripts, and the right-hand side panel shows genes within the “Function unknown” category. We highlighted enriched categories with an asterisk (* *p*-value < 0.05).

Figure S3 | Venn diagrams of putative post-transcriptionally regulated genes shared among different physiological states. **A**. Entities with proteins within the lower quintile of protein levels or not detected by our proteome survey whose mRNA levels are within the upper quintile (union set = 167). **B**. Entities within the lower quintile of protein levels and within the upper quintile of mRNA levels (union set = 64). **C**. Entities with proteins not detected by our proteome survey and within the upper quintile of mRNA levels (union set = 117). TP1: early exponential growth phase; TP2: mid-exponential growth phase; TP3: late exponential growth phase; TP4: stationary phase. All sets are available in File S3.

Figure S4 | Atlas section of putative post-transcriptionally regulated genes in the transition from TP1 to TP2. This section of the atlas shows genes having downregulated proteins and upregulated mRNAs (green cluster in Figure 2H) in the transition from the early exponential growth phase (TP1) to mid-exponential growth phase (TP2). The heatmap represents log_10_-transformed expression profile of proteins (a pseudocount was imputed for missing values), mRNAs (TPM+1), and ribosome-protected mRNA fragments (RPF; TPM+1). Heatmaps also represent the respective log_2_-transformed translational efficiency (TE) and ribosome occupancy (RO) for each time point. COG: clusters of orthologous genes; asRNAs: antisense RNA; TPS: transcript processing site; 2099: log_2_ fold change (LFC) of transcripts in the absence of RNase_2099C; CAI: codon adaptation index; TP3: late exponential growth phase; TP4: stationary phase.

Figure S5 | UpSet plot of putative post-transcriptionally regulated genes shared in different physiological state transitions. Entities being downregulated at the protein level and upregulated at the mRNA level (union set = 26). TP1: early exponential growth phase; TP2: mid-exponential growth phase; TP3: late exponential growth phase; TP4: stationary phase. All sets are available in File S6.

Figure S6 | Protein levels are associated with transcript GC content. The solid line illustrates the locally weighted smoothing (loess), and the shaded gray ribbon indicates its 95% confidence interval. A dashed line indicates the average GC content computed using the whole set of transcripts. Points follow a color gradient defined by the codon adaptation index (CAI). TP1: early exponential growth phase; TP2: mid-exponential growth phase; TP3: late exponential growth phase; TP4: stationary phase.

Figure S7 | VNG_0112H, a transposase encoded by the IS*H3B* element. Tracks show various features described on the left-hand side of the panel. Green tick marks represent transcript processing sites (TPS); red rectangles represent SmAP1 binding sites; a blue rectangle (reverse strand) represents the open reading frame for the transposase VNG_0112H; a green rectangle (reverse strand) represent the IS*H3B* element. Gray single-nucleotide resolution bar plots represent RNA-Seq and Ribo-Seq coverage; TP2: mid-exponential growth phase.

Figure S8 | Detected mobilization events. **A**. Detected insertions. **B**. Detected excisions. Observed events are the number of detected clusters for each type of mobilization. All the cluster types are represented, considering those classified as predominant, common, and rare. Bars are color-coded according to insertion sequence families.

Figure S9 | Protein-mRNA dynamics and various features of genes encoding gas vesicle biogenesis proteins. We represented the 14 genes comprising the *gvpDEFGHIJKLM* and *gvpACNO* operons in the context of their features. SmAP1 binding, antisense RNAs (asRNAs), and transcript processing sites (TPS) are enriched in this cluster (*p*-value = 2.4×10^−7^, 3×10^−3^, and 3.8×10^−2^, respectively). The heatmap represents log_10_-transformed expression profile of proteins (a pseudocount was imputed for missing values), mRNAs (TPM+1), and ribosome-protected mRNA fragments (RPF; TPM+1). Heatmaps also represent the respective log_2_-transformed translational efficiency (TE) and ribosome occupancy (RO) for each time point. COG: clusters of orthologous genes; 2099: log_2_ fold change (LFC) of transcripts in the absence of RNase_2099C; CAI: codon adaptation index; TP1: early exponential growth phase; TP2: mid-exponential growth phase; TP3: late exponential growth phase; TP4: stationary phase.

Figure S10 | *gvpACN* loci reveal differential patterns of Ribo-Seq signal. We present the three consecutive loci (VNG_7025-VNG_7027) comprising the *gvpACN* region (blue rectangles). The time point-wise Ribo-Seq and RNA-Seq normalized profiles are represented by gray bars. Red rectangles represent SmAP1 binding sites; green tick marks represent transcript processing sites (TPS); purple rectangles represent antisense RNAs. Each track was automatically scaled using the “Autoscale” feature of Integrative Genomics Viewer. We observe that pile-ups of Ribo-Seq emerge after the late exponential growth phase (TP3), indicating that the elongation phase of translation intensifies late on growth. Concurrently, we see SmAP1 binding sites either right before or spanning the region where the peaks emerge, indicating the role of this protein as a translational regulator. TP1: early exponential growth phase; TP2: mid-exponential growth phase; TP4: stationary phase.

Figure S11 | VNG_0042G, a TnpB encoded by the IS*H39* element from the IS*200*/IS*605* family subgroup IS*1341*. Tracks show various features described on the left-hand side of the panel. Green tick marks represent transcript processing sites (TPS); red rectangles represent SmAP1 binding sites; a purple rectangle (forward strand) represent an annotated antisense RNA; a blue rectangle (reverse strand) represents the open reading frame for TnpB; a green rectangle (reverse strand) represents the IS*H39* element. Gray single-nucleotide resolution bar plots represent RNA-Seq and Ribo-Seq coverage; TP2: mid-exponential growth phase.

Figure S12 | Growth curve of Δ*ura3* and Δ*ura3*Δ*smap1* strains. We conducted a growth curve experiment with three biological replicates for Δ*ura3* (blue lines) and Δ*ura3*Δ*smap1* (orange lines) strains. Line types depict each of the biological replicates.

## SUPPLEMENTAL FILES

File S1 | Atlas data. The non-redundant transcriptome locus tag dictionary, the normalized atlas data, and the non-normalized atlas data.

File S2 | Differentially expressed genes in the absence of RNase_2099C.

File S3 | Putative post-transcriptionally regulated genes (absolute abundance-based approach). Genes with patterns compatible with the post-transcriptional regulation hypothesis found by the abundance-based approach.

File S4 | Gene set enrichment analysis and comparison of features. Comparison of quantitative variables and enrichment tests for putative post-transcriptionally regulated gene sets found by the absolute abundance- and by the relative abundance-based approaches.

File S5 | Differential expression analysis of transcripts and proteins across the growth curve.

File S6 | Putative post-transcriptionally regulated genes (relative abundance-based approach). Genes with patterns compatible with the post-transcriptional regulation hypothesis found by the relative abundance-based approach.

File S7 | Atlas heatmap (expanded version). This file brings an expanded version of Figure 3. File S8 | Insertion sequence mobilization events detected by the long-read DNA-Seq experiment.

