## Supplemental Tables S1-6 and Figures S1-12 for "A genome-scale atlas reveals complex interplay of transcription and translation in an archaeon"

Affiliations:

| Annotation source | Reference |
|---|---|
| Third-Party Annotation section of GenBank (BK010829, BK010830, and BK010831). | (1) |
| NCBI Assembly RefSeq annotation for *H. salinarum* NRC-1 (ASM680v1) | (2) |
| NCBI Assembly RefSeq annotation for *H. salinarum* R1 (ASM6902v1) | (2) |
| Derived from original *H. salinarum* NRC-1 genome assembly and stored at Institute for Systems Biology SBEAMS. | In-house resource |

**Table S1 | Annotation sources for constructing the *Halobacterium salinarum* NRC-1 non-redundant transcriptome and a loci dictionary.**

| Pearson correlation coefficient comparison | 95% CI of Δ$R$ * | Significant difference |
|---|---|---|
| Nonoverlapping correlations based on dependent groups | | |
| $R_{TP1(A)}$ vs. $R_{TP2(B)}$ (0.6783 vs. 0.6869) | [-0.0195, 0.0019] | No |
| $R_{TP2(B)}$ vs. $R_{TP3(C)}$ (0.6833 vs. 0.5932) | [0.0702, 0.1112] | Yes |
| $R_{TP3(C)}$ vs. $R_{TP4(D)}$ (0.5724 vs. 0.4739) | [0.0767, 0.1213] | Yes |
| Overlapping correlations based on dependent groups | | |
| $R_{P\text{-}TP2\ m\text{-}TP1(E)}$ vs. $R_{TP2(B)}$ (0.6839 vs. 0.6826) | [-0.0079, 0.0106] | No |
| $R_{P\text{-}TP3\ m\text{-}TP2(F)}$ vs. $R_{TP3(C)}$ (0.6669 vs. 0.5704) | [0.0735, 0.1204] | Yes |
| $R_{P\text{-}TP4\ m\text{-}TP3(F)}$ vs. $R_{TP4(D)}$ (0.5666 vs. 0.4439) | [0.1015, 0.1450] | Yes |

**Table S2 | Comparison of Pearson correlation coefficient computed for protein and mRNA abundance throughout the growth curve.** We compared the coefficients using Zou's confidence interval method (3) implemented in the cocor package (4). Subscripts *P* and *m* refer to protein and mRNA levels for indicated time points. Uppercase letters (A-F) refer to panels in Figure 2. * Δ$R$ stands for the subtraction between the two coefficients (e.g., $R_{TP1(A)}$ - $R_{TP2(B)}$). A confidence interval (CI) of Δ$R$ spanning zero is not significant. Coefficients diverge slightly from those presented in the main text due to technical differences between the comparative approach and classic correlation method implementations. TP1: early exponential growth phase; TP2: mid-exponential growth phase; TP3: late exponential growth phase; TP4: stationary phase.

| IS name | Locus tag | Protein product | Family (Subgroup) | Transposition mechanism |
|---|---|---|---|---|
| Hasal_ISNpe8 | VNG_0056H | IS*H10*-type transposase | - | |
| HsIRS06 | VNG_6221H | IS*1341*-type transposase | - | |
| HsIRS12 | VNG_0028C | IS*H16*-type transposase | IS*630* | Cut-and-paste |
| HsIRS44 | VNG_6292C | IS*H14*-type transposase | IS*6* | Cointegrate |
| IS*H1* | VNG_0035H | IS*H9*-type transposase | IS*5* (IS*H1*) | - |
| IS*H2* | VNG_0210H | insertion element protein | IS*4* | Possibly cut-and-paste |
| IS*H3B* | VNG_0112H | IS*H3*-type transposase | IS*H3* | - |
| IS*H3C* | VNG_0051a | IS*H3*-type transposase | IS*H3* | - |
| IS*H4* | VNG_0918H | IS*H4*-type transposase | IS*1595* (IS*H4*) | - |
| IS*H7* | VNG_7005 | IS*H7*-type transposase | ISNCY | - |
| IS*H8A* | VNG_6038H | IS*H8*-type transposase | IS*4* (IS*H8*) | Cut-and-paste |
| IS*H8B* | VNG_0059H | IS*H8*-type transposase | IS*4* (IS*H8*) | Cut-and-paste |
| IS*H8C* | VNG_7119 | IS*H8*-type transposase | IS*4* (IS*H8*) | Cut-and-paste |
| IS*H8D* | VNG_0052H | IS*H8*-type transposase | IS*4* (IS*H8*) | Cut-and-paste |
| IS*H8E* | VNG_1653H | IS*H8*-type transposase | IS*4* (IS*H8*) | Cut-and-paste |
| IS*H9* | VNG_7034 | IS*H9*-type transposase | IS*5* (IS*H1*) | - |
| IS*H10* | VNG_0021H | IS*H10*-type transposase | IS*66* (ISBst12) | - |
| IS*H11* | VNG_0213H | IS*H11*-type transposase | IS*5* | - |
| IS*H12* | VNG_0043H, VNG_0044H | IS*200*-type transposase, IS*1341*-type transposase | IS*200*/IS*605* (IS*605*) | Peel-and-paste |
| IS*H22* | VNG_0285C, VNG_0286C | IS*200*-type transposase, IS*1341*-type transposase | IS*200*/IS*605* (IS*605*) | Peel-and-paste |
| IS*H29* | VNG_6430C | IS*H14*-type transposase | IS*6* | Cointegrate |
| IS*H32* | VNG_6148H | IS*H8*-type transposase | IS*4* (IS*H8*) | Cut-and-paste |

| IS name | Locus tag | Protein product | Family (Subgroup) | Transposition mechanism |
|---|---|---|---|---|
| IS*H34* | VNG_2652H, VNG_2653C | IS*1341*-type transposase, IS*200*-type transposase | IS*200*/IS*605* (IS*605*) | Peel-and-paste |
| IS*H35* | VNG_6181H, VNG_6182H | IS*1341*-type transposase, IS*200*-type transposase | IS*200*/IS*605* (IS*605*) | Peel-and-paste |
| IS*H37* | VNG_0013C | IS*1341*-type transposase | IS*200*/IS*605* (IS*1341*) | Peel-and-paste |
| IS*H38* | VNG_0026C | IS*1341*-type transposase | IS*200*/IS*605* (IS*1341*) | Peel-and-paste |
| IS*H39* | VNG_0042G | IS*1341*-type transposase | IS*200*/IS*605* (IS*1341*) | Peel-and-paste |
| IS*H40* | VNG_6361G | IS*1341*-type transposase | IS*200*/IS*605* (IS*605*) | Peel-and-paste |

**Table S3 | The non-redundant set of insertion sequences in *Halobacterium salinarum* NRC-1.** We obtained insertion sequence families from ISfinder (5) and ISsaga (6), and the transposition mechanisms from (7).

| Strain | Insertions (I) [a] | Excisions (E) [b] | Reads [c] | I + E Normalized [d] |
|---|---|---|---|---|
| Δ*ura3*Δ*smap1* A | 43 | 7 | 20,530 | 50.00 |
| Δ*ura3*Δ*smap1* B | 34 | 5 | 20,142 | 39.75 |
| Δ*ura3*Δ*smap1* C | 44 | 5 | 19,858 | 50.66 |
| Δ*ura3* A | 39 | 5 | 8,871 | 101.83 |
| Δ*ura3* B | 21 | 4 | 9,966 | 51.50 |
| Δ*ura3* C | 24 | 4 | 5,216 | 110.21 |

**Table S4 | Summary of the transposition detection assay.** [a] Number of identified insertion clusters. [b] Number of identified excision clusters. [c] Number of reads aligned to the reference genome. [d] Sum of insertion and excision clusters normalized by the library with the highest number of aligned reads.

| Application | Primer name | Sequence |
|---|---|---|
| SmAP1 null mutant construction and genotyping of genomic DNA samples for long-read sequencing | vng1496g_a | 5’-GCGACGGTGTACGCAGTGAGC |
| | vng1496g_b | 5’-ACCGCCGGTGGTGGCATCCAT |
| | vng1496g_c | 5’-GATGCCACCACCGGCGGTCCATGACTGGCGCAGGAACCCC |
| | vng1496g_d | 5’-AGGGCGTACAGCGCGTAGTAC |
| Genotyping of genomic DNA samples for long-read sequencing | vng1673g.a-ura3-ISB | 5’-CGGCACAGGAGGGCGTGCTGG |
| | vng1673g.d-ura3-ISB | 5’-GGTCCCGGGACCGCCCCCCGC |
| DNase treatment test of co-immunoprecipitated samples for RIP-Seq | 19-fwd | 5’-AACCACCGTGACCCGGAAGC |
| | 20-rev | 5’-GCCGATCCGATGGGAGTGACT |
| | 63-fwd | 5’-GTGAGGGACGAAAGCTAGGG |
| | 64-rev | 5’-AAGGGTACTGCTGGCAATTA |

**Table S5 | List of primers used in this study.**

| Source | Description | Uniform resource locator (URL) |
|---|---|---|
| In-house | Data integration, analysis, figures, supplemental material output, ShinyApp development. | https://github.com/alanlorenzetti/**halo_atlas** |
| | Generation of RNA-Seq signal visualization. | https://github.com/alanlorenzetti/**frtc** |
| | DNA-Seq structural variant analysis. | https://github.com/alanlorenzetti/**transpositionAnalysis_v2** |
| | RIP-Seq analysis. | https://github.com/alanlorenzetti/**ripper** |
| | RNA-Seq and Ribo-Seq quantitation using a non-redundant transcriptome. | https://github.com/alanlorenzetti/**runKallisto** |
| | *H. salinarum* NRC-1 non-redundant transcriptome. | https://github.com/alanlorenzetti/**halo_nr_tx** |
| | *H. salinarum* NRC-1 Atlas web portal implementation. | https://github.com/baliga-lab/**halodata** |
| Third-party | Computation of codon adaptation index (CAI). | https://github.com/BioinfoHR/**coRdon** |
| | Getting status for Oxford Nanopore Technologies DNA-Seq data. | https://github.com/rrwick/**Filtlong** |
| | Trimming of Oxford Nanopore Technologies DNA-Seq data. | https://github.com/rrwick/**Porechop** |

**Table S6 | In-house and third-party GitHub repositories cited in this study.**

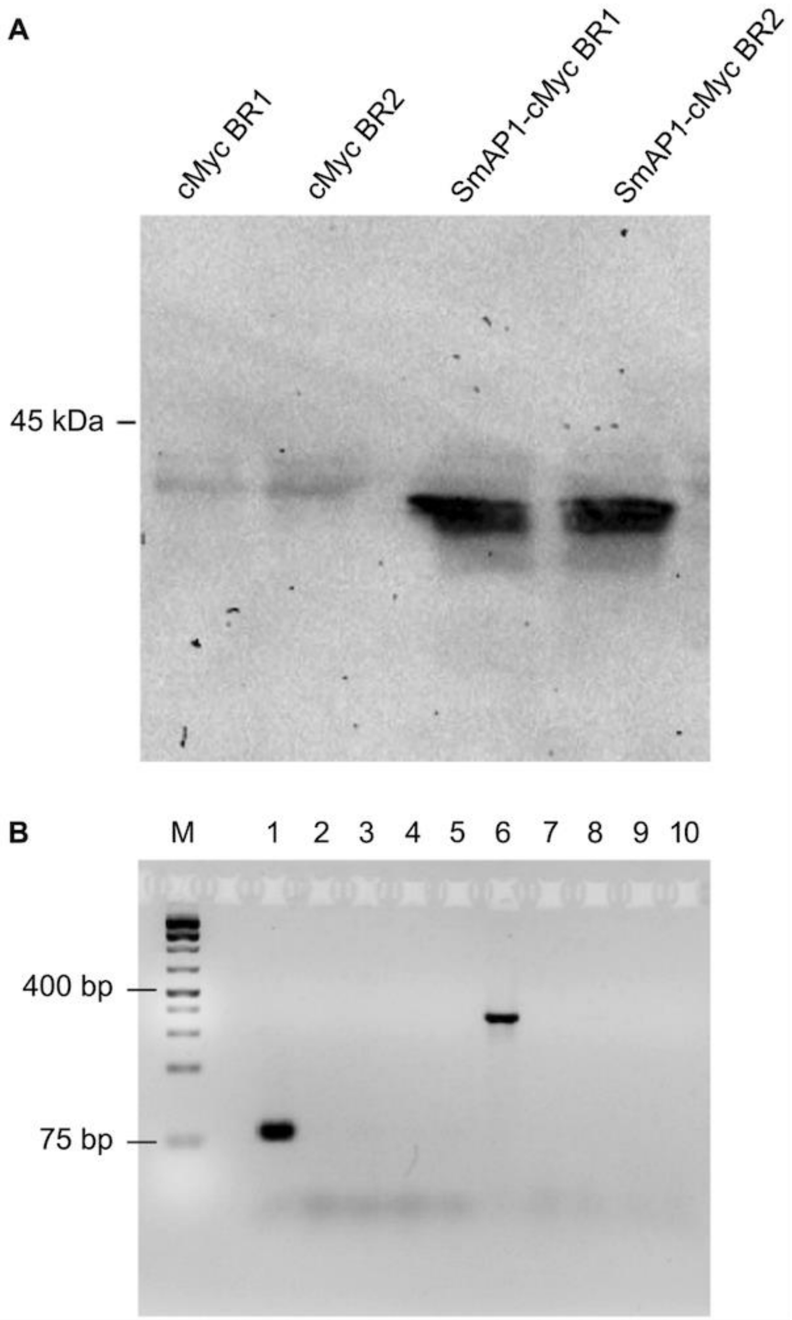

**Figure S1 | Quality assurance of co-immunoprecipitated samples. A.** Western blot of samples extracted from strains expressing plasmids for cMyc and cMyc-tagged SmAP1 (see lane titles for labels). The expected molecular weight of the cMyc-tagged SmAP1 complex is 37 kDa. BR: Biological replicate. **B.** Polymerase Chain Reaction (PCR) of RNA-purified samples treated with DNase. M: Ladder; 1: Positive control (genomic DNA amplified using 19-fwd and 20-rev primers with a predicted amplicon size of 85 bp); 2-5: cMyc BR1, cMyc BR2, SmAP1-cMyc BR1, and SmAP1-cMyc BR2 (amplified using 19-fwd and 20-rev primers); 6: Positive control (genomic DNA amplified using 63-fwd and 64-rev primers with a predicted amplicon size of 450 bp). 7-10: cMyc BR1, cMyc BR2, SmAP1-cMyc BR1, and SmAP1-cMyc BR2 (amplified using 63-fwd and 64-rev primers).

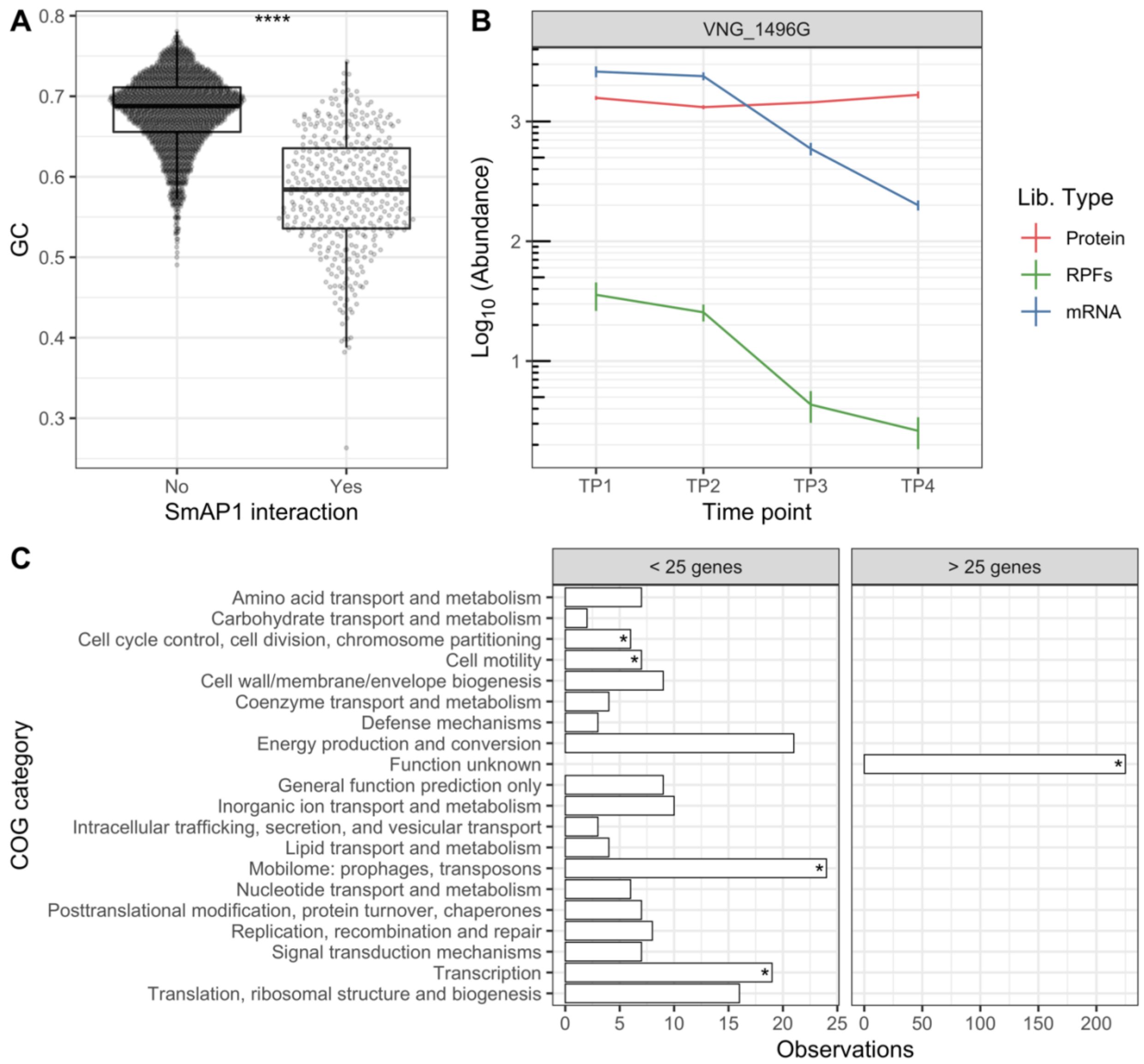

**Figure S2 | SmAP1 features. A.** SmAP1 binding is conditioned to the GC content of transcripts. The reduced GC content of transcripts is a property influencing SmAP1 binding. We compared medians using the Mann–Whitney U test. **** $p$-value ≤ $10^{-4}$. **B.** Time course view of protein, ribosome-protected mRNA fragments (RPF; TPM+1), and mRNA levels (TPM+1). Vertical bars represent the standard error computed using at least six replicates for proteins and three replicates for mRNA and RPF. **C.** Functional categories of transcripts bound to SmAP1. The panel shows how many genes have transcripts bound to SmAP1, considering each category of COG (clusters of orthologous genes). The left-hand side panel shows categories with no more than 25 genes with SmAP1-bound transcripts, and the right-hand side panel shows genes within the "Function unknown" category. We highlighted enriched categories with an asterisk (* $p$-value < 0.05).

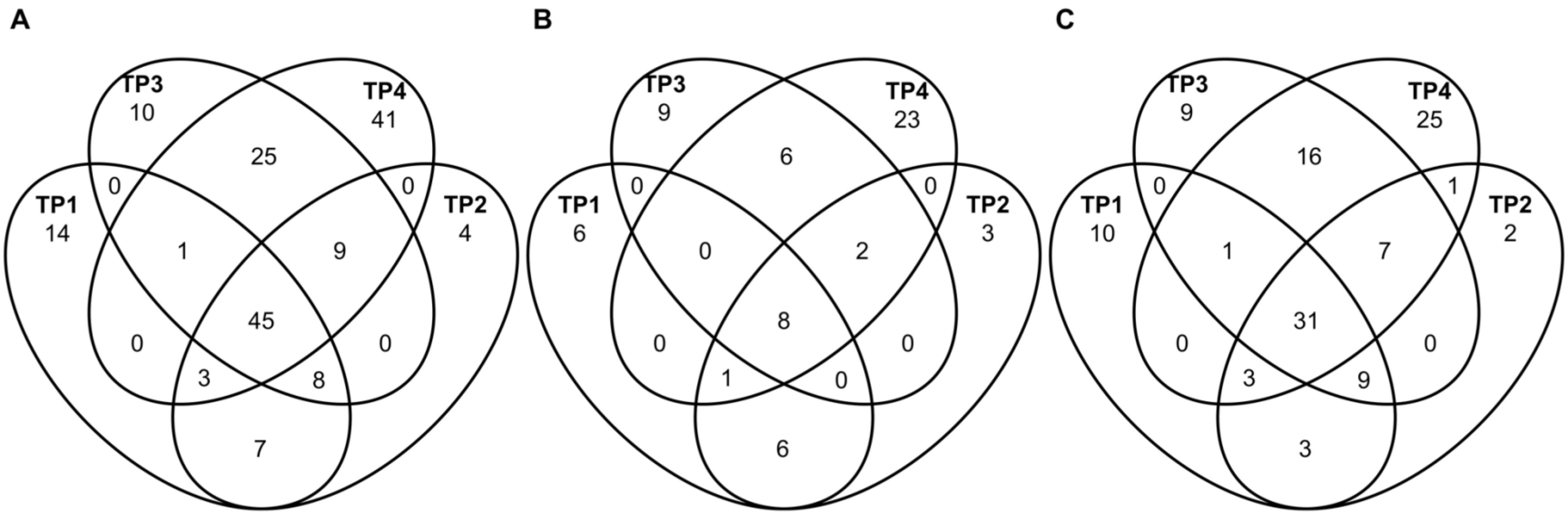

**Figure S3 | Venn diagrams of putative post-transcriptionally regulated genes shared among different physiological states. A.** Entities with proteins within the lower quintile of protein levels or not detected by our proteome survey whose mRNA levels are within the upper quintile (union set = 167). **B.** Entities within the lower quintile of protein levels and within the upper quintile of mRNA levels (union set = 64). **C.** Entities with proteins not detected by our proteome survey and within the upper quintile of mRNA levels (union set = 117). TP1: early exponential growth phase; TP2: mid-exponential growth phase; TP3: late exponential growth phase; TP4: stationary phase. All sets are available in File S3.

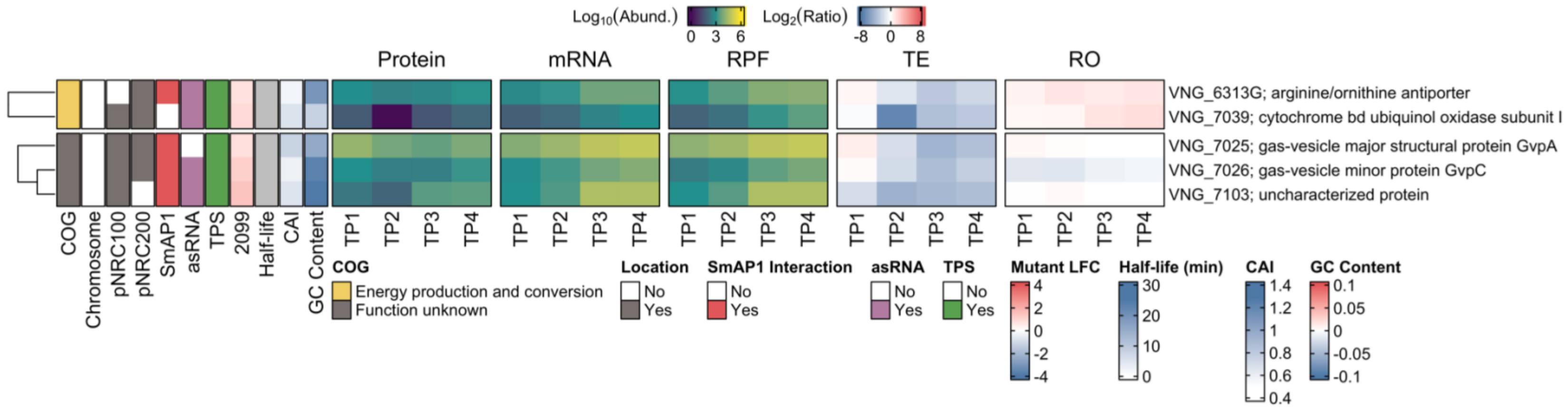

**Figure S4 | Atlas section of putative post-transcriptionally regulated genes in the transition from TP1 to TP2**. This section of the atlas shows genes having downregulated proteins and upregulated mRNAs (green cluster in Figure 2H) in the transition from the early exponential growth phase (TP1) to mid-exponential growth phase (TP2). The heatmap represents $log_{10}$-transformed expression profile of proteins (a pseudocount was imputed for missing values), mRNAs (TPM+1), and ribosome-protected mRNA fragments (RPF; TPM+1). Heatmaps also represent the respective $log_2$-transformed translational efficiency (TE) and ribosome occupancy (RO) for each time point. COG: clusters of orthologous genes; asRNAs: antisense RNA; TPS: transcript processing site; 2099: $log_2$ fold change (LFC) of transcripts in the absence of RNase_2099C; CAI: codon adaptation index; TP3: late exponential growth phase; TP4: stationary phase.

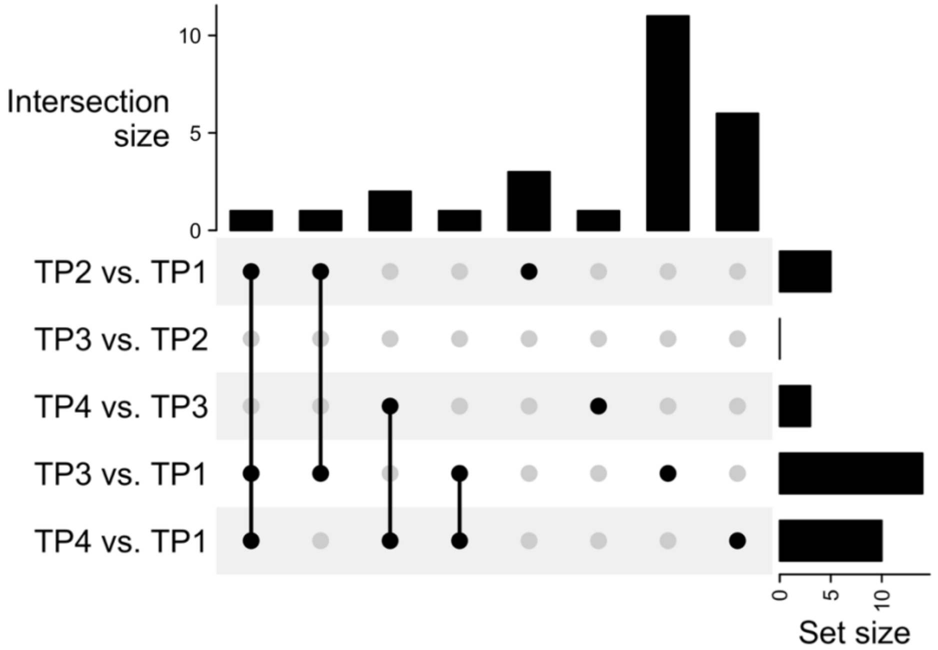

**Figure S5 | UpSet plot of putative post-transcriptionally regulated genes shared in different physiological state transitions**. Entities being downregulated at the protein level and upregulated at the mRNA level (union set = 26). TP1: early exponential growth phase; TP2: mid-exponential growth phase; TP3: late exponential growth phase; TP4: stationary phase. All sets are available in File S6.

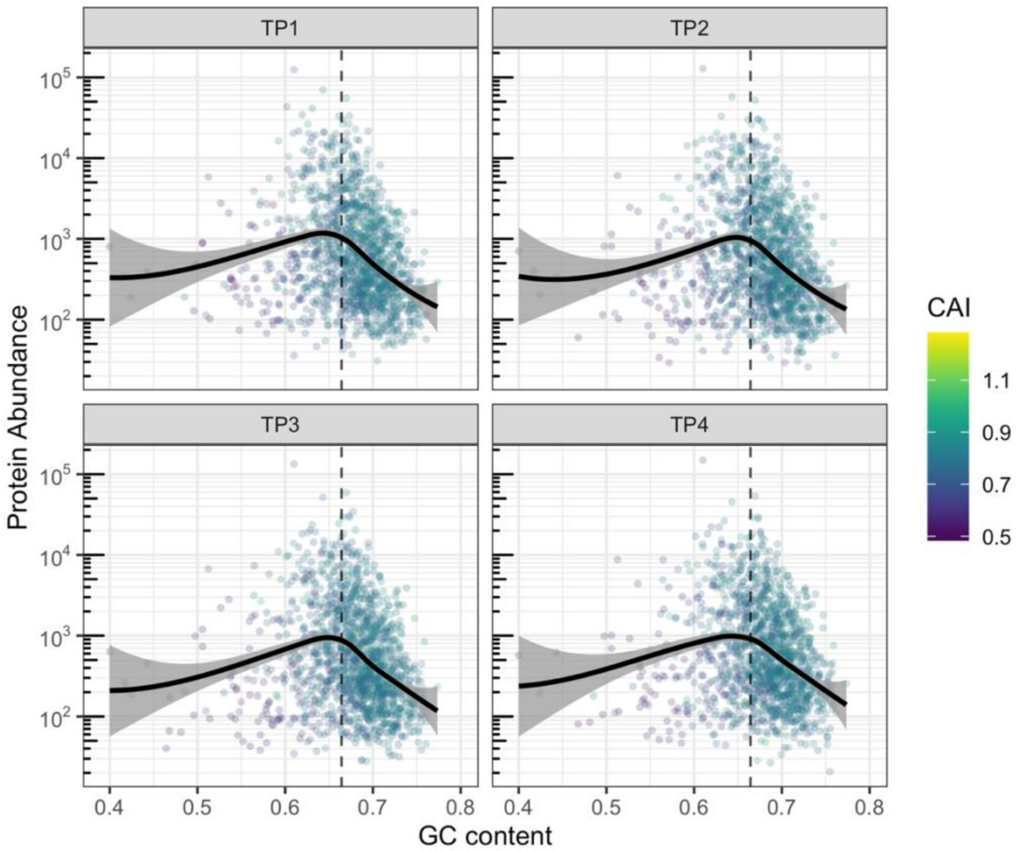

**Figure S6 | Protein levels are associated with transcript GC content.** The solid line illustrates the locally weighted smoothing (loess), and the shaded gray ribbon indicates its 95% confidence interval. A dashed line indicates the average GC content computed using the whole set of transcripts. Points follow a color gradient defined by the codon adaptation index (CAI). TP1: early exponential growth phase; TP2: mid-exponential growth phase; TP3: late exponential growth phase; TP4: stationary phase.

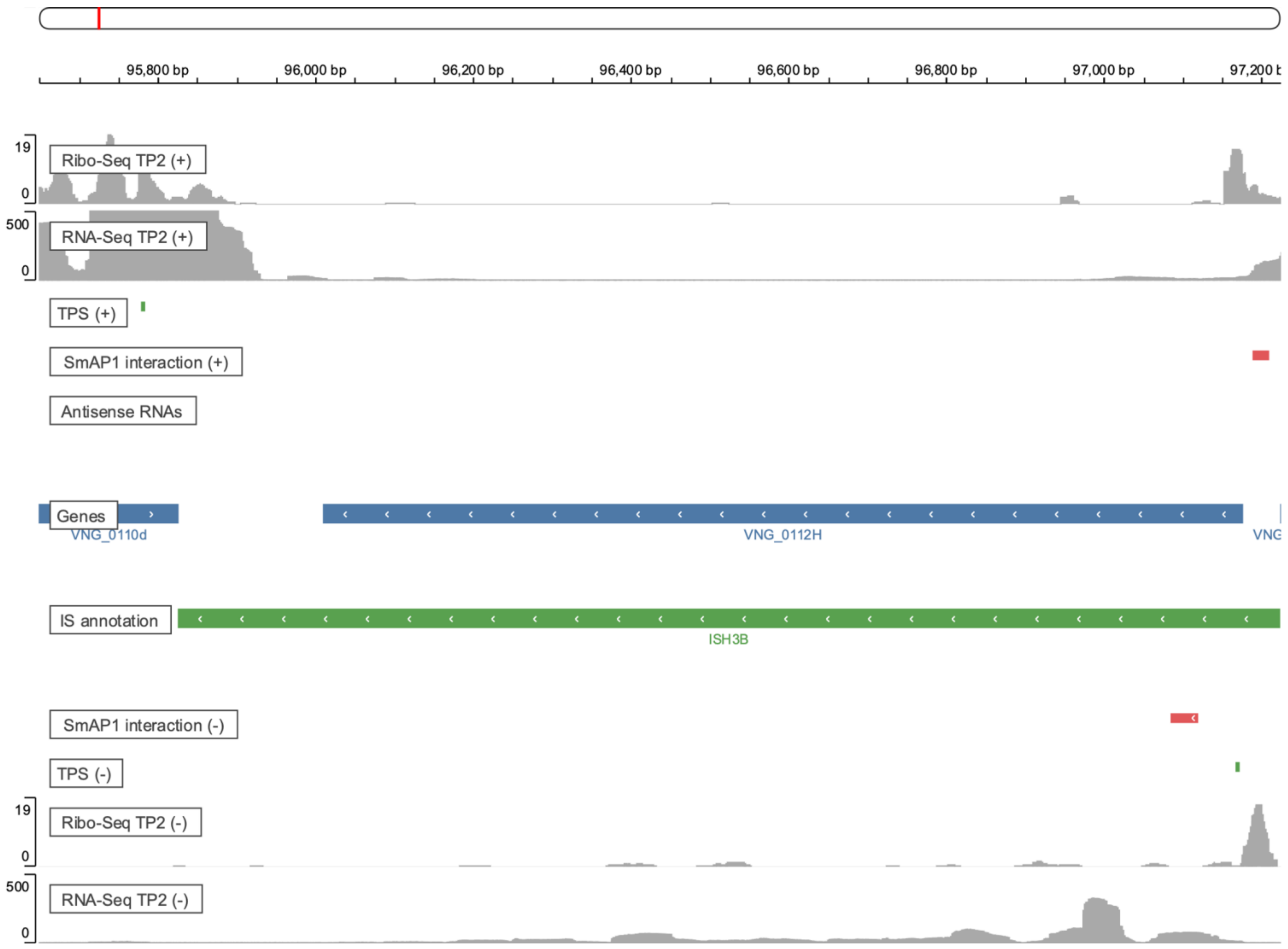

**Figure S7 | VNG_0112H, a transposase encoded by the IS*H3B* element.** Tracks show various features described on the left-hand side of the panel. Green tick marks represent transcript processing sites (TPS); red rectangles represent SmAP1 binding sites; a blue rectangle (reverse strand) represents the open reading frame for the transposase VNG_0112H; a green rectangle (reverse strand) represent the IS*H3B* element. Gray single-nucleotide resolution bar plots represent RNA-Seq and Ribo-Seq coverage; TP2: mid-exponential growth phase.

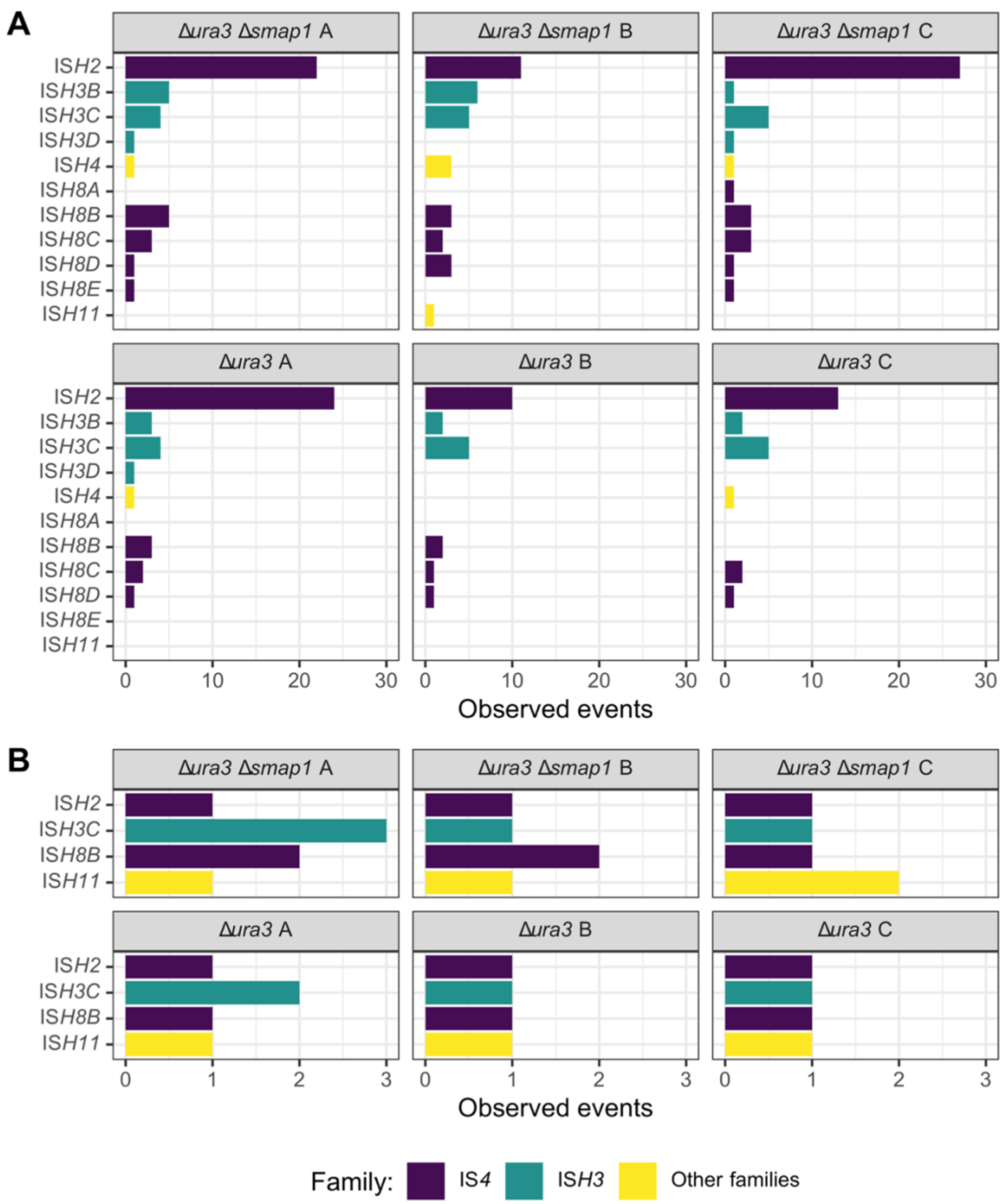

**Figure S8 | Detected mobilization events. A**. Detected insertions. **B.** Detected excisions. Observed events are the number of detected clusters for each type of mobilization. All the cluster types are represented, considering those classified as predominant, common, and rare. Bars are color-coded according to insertion sequence families.

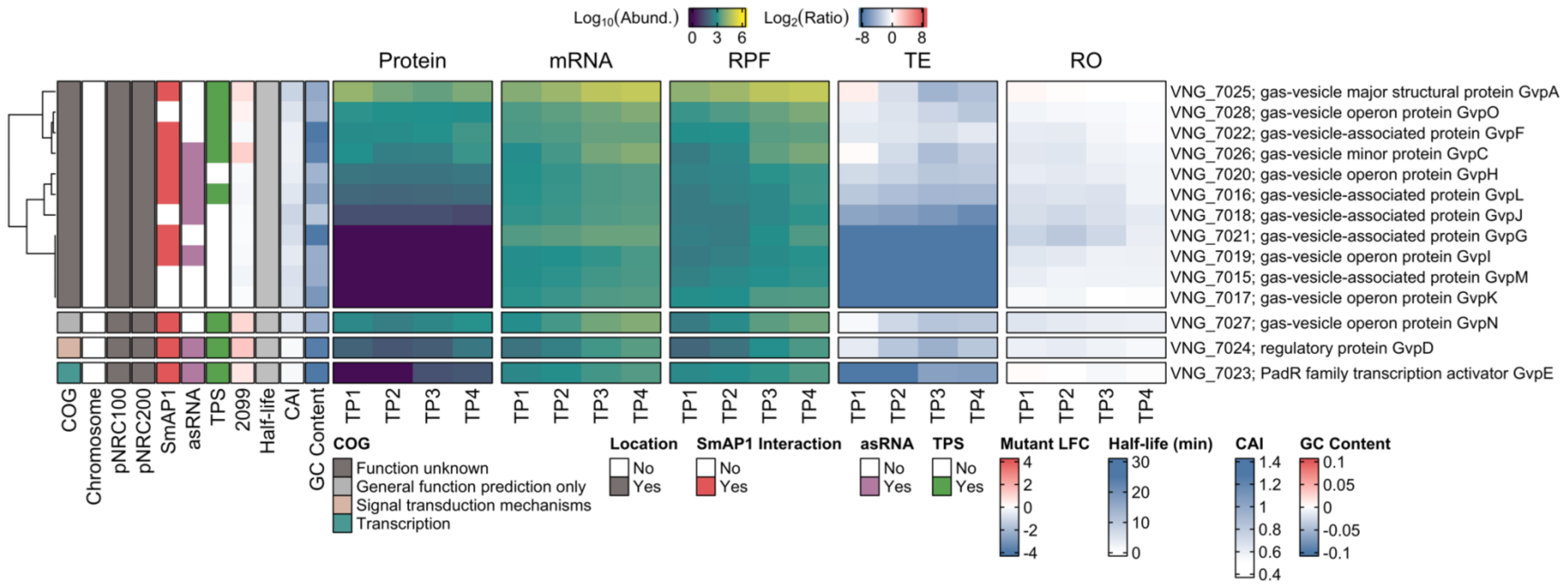

**Figure S9 | Protein-mRNA dynamics and various features of genes encoding gas vesicle biogenesis proteins.** We represented the 14 genes comprising the *gvpDEFGHIJKLM* and *gvpACNO* operons in the context of their features. SmAP1 binding, antisense RNAs (asRNAs), and transcript processing sites (TPS) are enriched in this cluster (*p*-value = $2.4 \times 10^{-7}$, $3 \times 10^{-3}$, and $3.8 \times 10^{-2}$, respectively). The heatmap represents $\log_{10}$-transformed expression profile of proteins (a pseudocount was imputed for missing values), mRNAs (TPM+1), and ribosome-protected mRNA fragments (RPF; TPM+1). Heatmaps also represent the respective $\log_2$-transformed translational efficiency (TE) and ribosome occupancy (RO) for each time point. COG: clusters of orthologous genes; 2099: $\log_2$ fold change (LFC) of transcripts in the absence of RNase_2099C; CAI: codon adaptation index; TP1: early exponential growth phase; TP2: mid-exponential growth phase; TP3: late exponential growth phase; TP4: stationary phase.

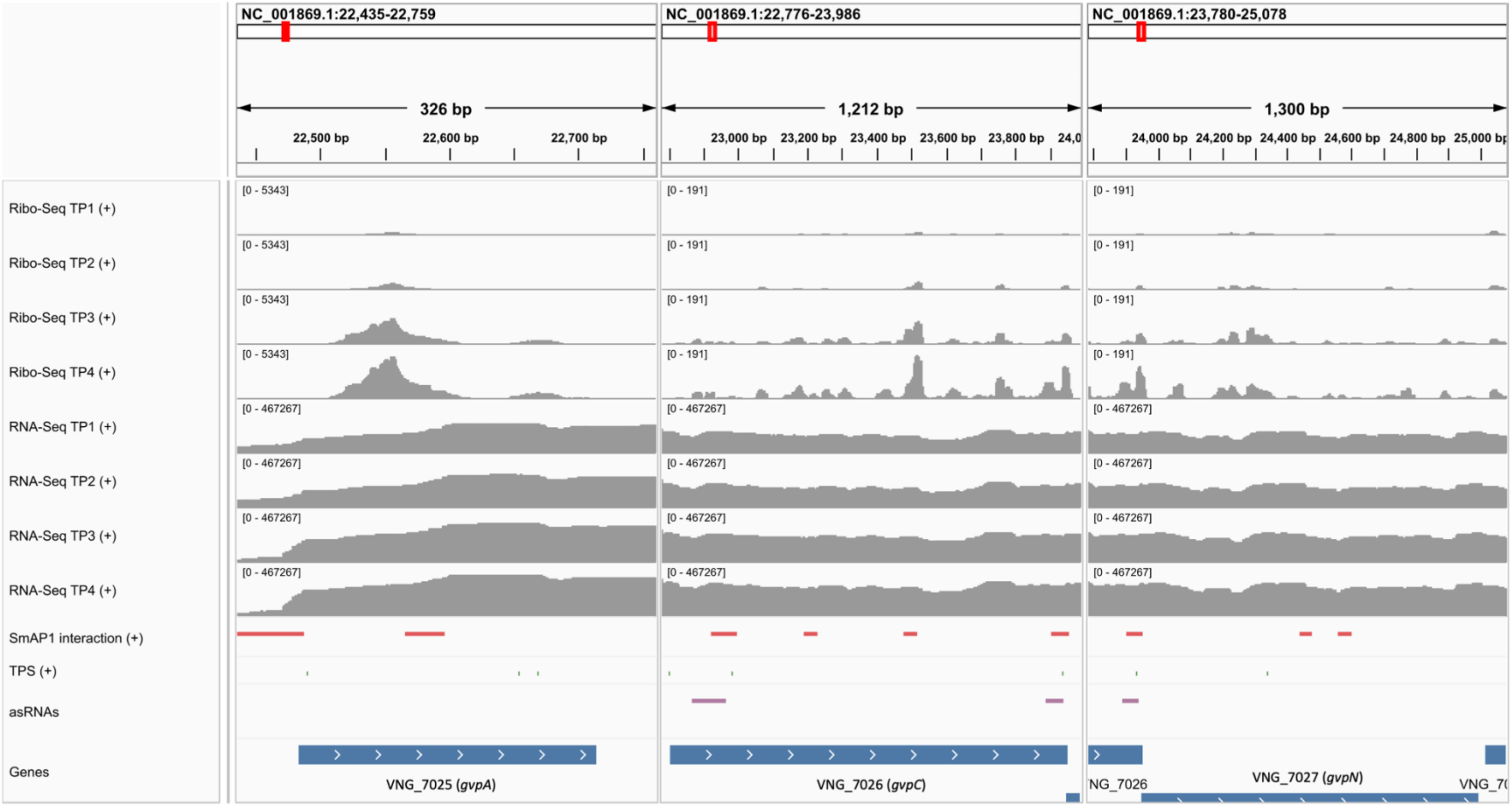

**Figure S10 | *gvpACN* loci reveal differential patterns of Ribo-Seq signal.** We present the three consecutive loci (VNG_7025-VNG_7027) comprising the *gvpACN* region (blue rectangles). The time point-wise Ribo-Seq and RNA-Seq normalized profiles are represented by gray bars. Red rectangles represent SmAP1 binding sites; green tick marks represent transcript processing sites (TPS); purple rectangles represent antisense RNAs. Each track was automatically scaled using the "Autoscale" feature of Integrative Genomics Viewer (8). We observe that pile-ups of Ribo-Seq emerge after the late exponential growth phase (TP3), indicating that the elongation phase of translation intensifies late on growth. Concurrently, we see SmAP1 binding sites either right before or spanning the region where the peaks emerge, indicating the role of this protein as a translational regulator. TP1: early exponential growth phase; TP2: mid-exponential growth phase; TP4: stationary phase.

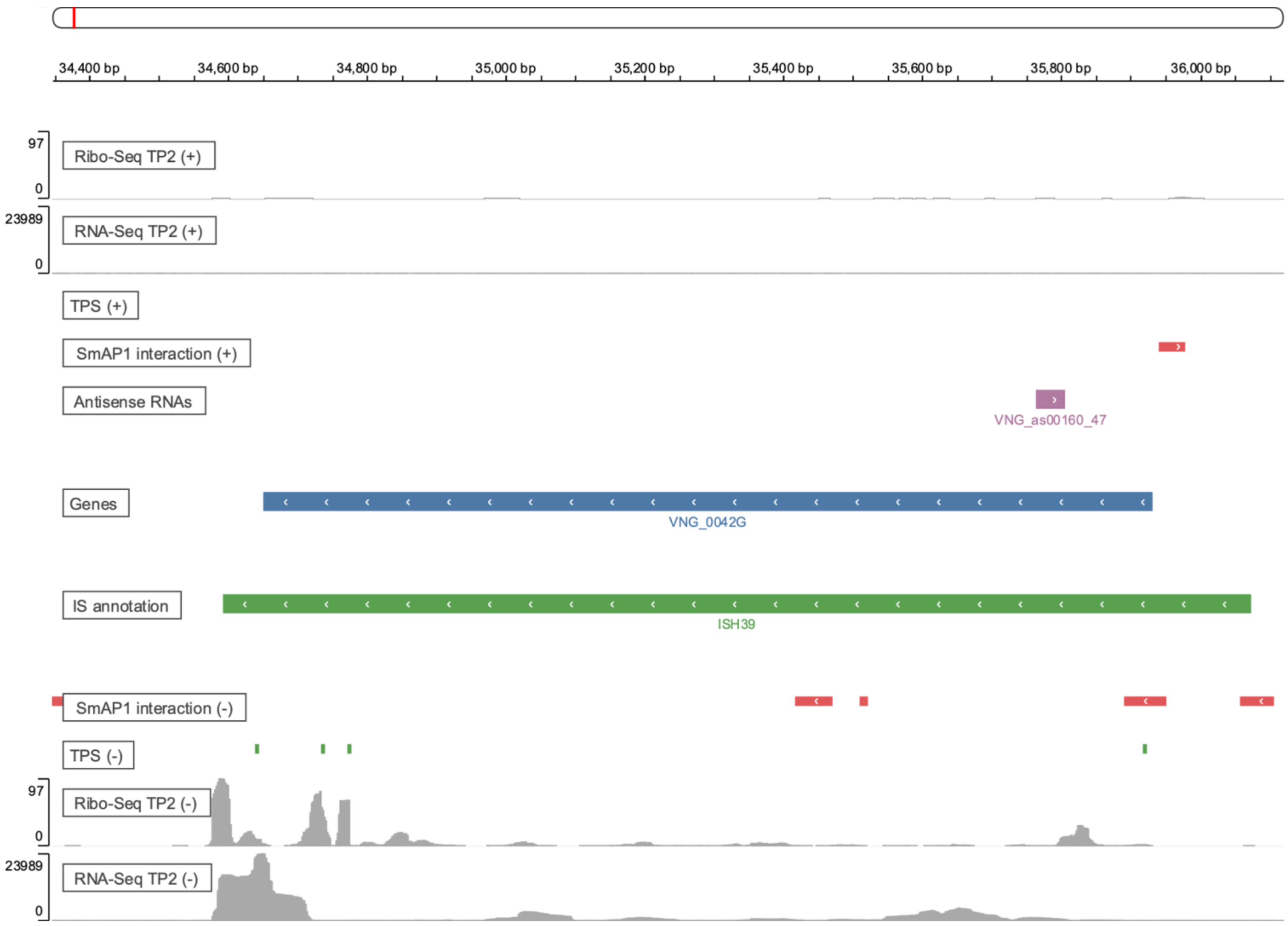

**Figure S11 | VNG_0042G, a TnpB encoded by the IS*H39* element from the IS*200*/IS*605* family subgroup IS*1341***. Tracks show various features described on the left-hand side of the panel. Green tick marks represent transcript processing sites (TPS); red rectangles represent SmAP1 binding sites; a purple rectangle (forward strand) represent an annotated antisense RNA; a blue rectangle (reverse strand) represents the open reading frame for TnpB; a green rectangle (reverse strand) represents the IS*H39* element. Gray single-nucleotide resolution bar plots represent RNA-Seq and Ribo-Seq coverage; TP2: mid-exponential growth phase.

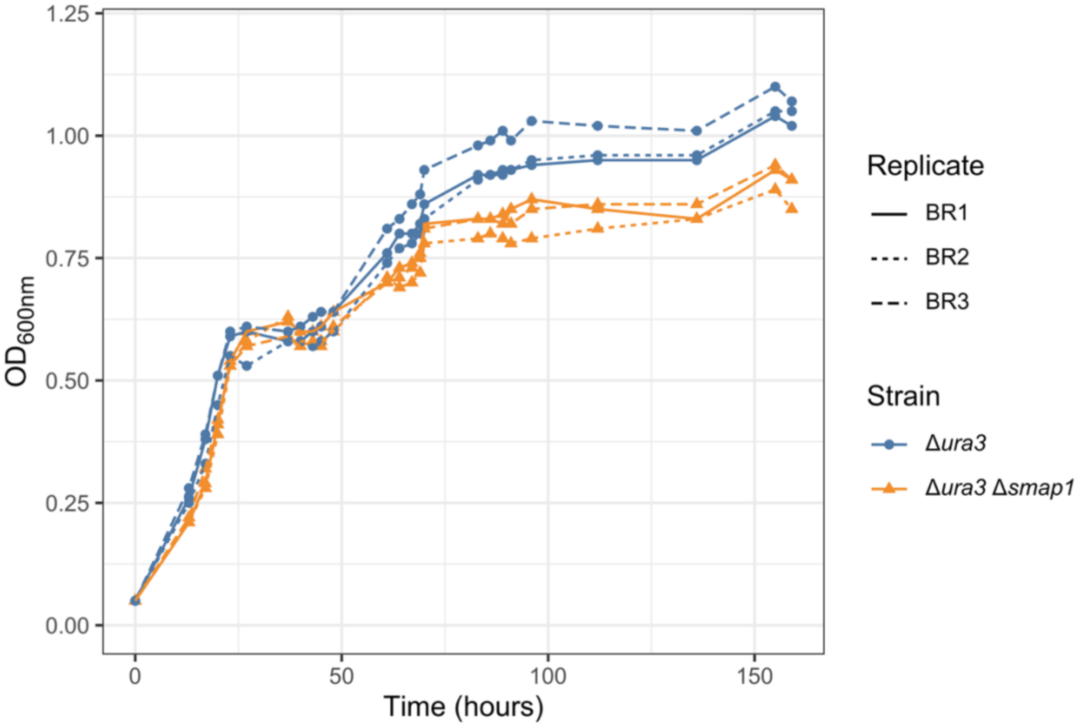

**Figure S12 | Growth curve of Δ*ura3* and Δ*ura3*Δ*smap1* strains.** We conducted a growth curve experiment with three biological replicates for Δ*ura3* (blue lines) and Δ*ura3*Δ*smap1* (orange lines) strains. Line types depict each of the biological replicates.
