## Supplementary figures and images for "A genome-scale atlas reveals complex interplay of transcription and translation in an archaeon"

### File S7

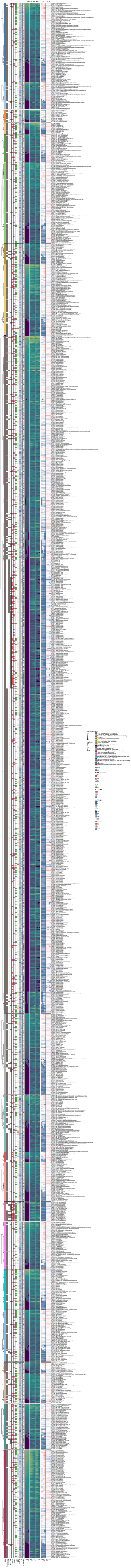
